# Sense organ formation and identity are controlled by divergent mechanisms in insects

**DOI:** 10.1101/2020.09.04.281865

**Authors:** Marleen Klann, Magdalena Ines Schacht, Matthew Alan Benton, Angelika Stollewerk

## Abstract

Insects and other arthropods utilise external sensory structures for mechanosensory, olfactory and gustatory reception. These sense organs have characteristic shapes related to their function, and in many cases are distributed in a fixed pattern so that they are identifiable individually. In *Drosophila melanogaster*, the identity of sense organs is regulated by specific combinations of transcription factors. In other arthropods, however, sense organ subtypes cannot be linked to the same code of gene expression. This raises the questions of how sense organ diversity has evolved in arthropods, and if the *D. melanogaster* subtype identity principle is representative for insects. To address these questions, we analyse sense organ development in another insect, the flour beetle *Tribolium castaneum.* We show that in contrast to *D. melanogaster*, *T. castaneum* sense organs cannot be categorised based on their requirement for individual or combinations of the conserved sense organ transcription factors such as *cut* and *pox-neuro* and members of the Achaete-Scute (*Tc ASH*, *Tc asense*) and Atonal family (*Tc atonal*, *Tc cato*, *Tc amos*). Rather, these genes are required for the specification of sense organ precursors and the development and differentiation of sensory cell types in diverse external sensilla. Based on our findings and past research, we present an evolutionary scenario suggesting that sensory organs have diversified from a default state through subsequent recruitment of sensory genes to the different sense organ specification processes. A specific role for genes in subtype identity has evolved as a secondary effect of the function of these genes in individual or subsets of sense organs, which can largely not be aligned with morphological or functional categories.

## Introduction

In arthropods, external sense organs function at the interface of the environment and the organism (Barth, 2001; Dangles et al., 2009; Hansson and Stensmyr, 2011, Stevens, 2013). Different types (and subtypes) of sense organs exist, all of which can generally be found across the arthropod body. However, some subtypes are clustered on specific appendages that are primarily used for specific behaviours, such as gustatory receptors on mouthparts or olfactory receptors on insect antennae (Shanbhag et al., 1999; Dahanukar et al., 2005; Joseph and Carlson, 2015). External sense organs show a great variety of habitat- and behaviour-adapted forms and functions, ranging from the simple mechanosensory bristles of flies to the complex cuticular structures of crustacean feeding setae (Chapman, 2013; Klann and Stollewerk, 2017). This diversity raises the question of how the different shapes and functions have emerged in arthropods and which molecular mechanisms have facilitated their evolution.

Although there is no uniform classification of sense organs in arthropods, external and internal sense organs are generally distinguished from one another (Hartenstein, 2005; Chapman, 2013). In our present study, we focus on external sense organs in insects. There are at least five morphological categories of external sense organs described across insect species: chaetoid, trichoid, basiconic, campaniform and coeloconic sensilla (Snodgrass, 1926; Stocker, 1994; Hartenstein, 2005; Chapman, 2013). These can be further subdivided and assigned functions based on additional characteristics (Table 1). For example, aporous trichoid sensilla are mechanosensory organs, while multiporous trichoid sensilla function as olfactory receptors (Hartenstein, 2005).

**Table 1.** Characteristics of external sensilla of insects.

| Sense organ | Structure | Additional characteristics | Function |
| --- | --- | --- | --- |
| **Chaetoid sensillum** | - bristle-like<br>- movably<br>- articulated in wide socket | - uniporous (pore at tip)<br>- steep angle | Contact chemosensory organ<sup>*1</sup> |
|  |  | - aporous<br>- flat angle | Mechanosensory organ<sup>*1</sup> |
| **Trichoid sensillum** | - hair-like<br>- articulated in tight socket | - uniporous (pore at tip)<br>- steep angle | Contact chemosensory organ<sup>*2</sup> |
|  |  | - multiporous<br>- curved hair | Olfactory sense organ<sup>*2</sup> |
|  |  | - aporous | Mechanosensory organ<sup>*2</sup> |
| **Basiconic sensillum** | - pegs<br>- cones | - blunt tip<br>- sharp or curved<br>- flat angle when mechanosensory | Mechano- & chemosensory organ<sup>*1,2</sup> |
| **Campaniform sensillum** | - dome with collar | - variations in collar size<br>- dome shape | Proprioception |
| **Coeloconic sensillum** | - pit with cone inside |  | Hygro/thermo-sensory |

<sup>\*1</sup> Ploomi et al., 2003; <sup>\*2</sup> Ryan and Behan, 1973

Arthropod sense organs arise from epithelial sensory organ progenitor cells (SOPs), which give rise to 4-5 different cell types (neurons, glia, sheath cells and cells generating the cuticular structure, e.g. hair, socket) (Hallberg and Hansson, 1999; Hartenstein, 2005; Chapman, 2013). In *Drosophila melanogaster*, 5 bHLH transcription factors determine which sense organ subtypes are generated by the SOPs (Hartenstein, 2005). However, the molecular subdivision is not in line with the 5 morphological categories (Table 1) (Ploomi et al., 2003; Ryan, 1973); rather, it classifies the physiological function of sense organs.

Members of the Achaete-scute family (*achaete* (*ac*), *scute* (*sc*), and *asense* (*ase*)) determine external mechanosensory and gustatory sense organs, while Atonal family members (*atonal* (*ato*) and *absent MD neurons and olfactory sensilla* (*amos*)) specify (external) olfactory and internal mechanosensory organs (Jarman et al., 1993a; Bertrand et al., 2002; Lai and Orgogozo, 2004; Hartenstein, 2005; Muang and Jarman, 2007).

These 5 transcription factors are required both for SOP formation as well as their subtype identity. Additional transcription factors, *cut (ct)*, *pox-neuro (poxn)*, *cousin of atonal* (*cato)*, and *target of pox-neuro* (*tap*), are expressed downstream of these transcription factors in subsets of sense organs and, when mutated, partially change subtype identity (Nottebohm et al., 1994; Gautier et al., 1997; Jarman and Ahmed, 1998; Ledent et al., 1998; Goulding et al., 2000a, 2000b; Brewster et al., 2001; Skaer et al., 2002; zur Lage et al.; 2003; Hartenstein, 2005; Lai et al., 2013). For example, mutations in *ct*, which is expressed in external mechanosensory and gustatory sensilla, lead to transformations of these sensilla into chordotonal organs (Bodmer et al., 1987). Another set of genes (e.g. *asense* (*ase*), *prospero* (*pros*) and *snail* (*sna*)) is required for the development of the different cell types within the sense organs (González et al., 1989; Doe et al., 1991; Dominguez and Campuzano, 1993; Brand et al., 1993; Jarman et al., 1993b; Ip et al., 1994; Hartenstein, 2005; Ayyar et al., 2010; Johnson et al., 2019).

Taken together, the *D. melanogaster* data show that a combinatorial code of transcription factors seems to determine sense organ subtype identity. This raises the question how this code has evolved and how, or if, it is used in other arthropod taxa or even in other insect species. Thus far, only a few studies on sense organ development in arthropods other than *D. melanogaster* exist. These studies have shown that genes known to be involved in sense organ subtype specification and cell type determination within the SOP lineage in *D. melanogaster* (*ASH*, *ato*, *ase*, *sna*, *pros*, *Notch*, *Numb*) are expressed during sense organ development. However, these genes seem to perform different/additional roles in different species (e.g. Gold et al., 2009). For example, *ato* shows a conserved expression in chemosensory organs of the crustacean *Daphnia magna* (Klann and Stollewerk, 2017), the myriapod *Glomeris marginata* (Pioro and Stollewerk, 2006) and the insect *D. melanogaster* but, in addition, is co-expressed with *ASH* in various types of sense organs in the crustacean, including mechanosensory sensilla (Klann and Stollewerk, 2017). Does this suggest that different codes for sense organ subtype specification have evolved in insects and the closely related crustaceans? Or is the *D. melanogaster* code not even representative for insects? In order to address this question, we analyse here the expression and function of sensory genes in the development of different categories of sensilla in the flour beetle *Tribolium castaneum*.

## Results

### Distribution of external sensilla in the first larval stage

In order to analyse the molecular mechanisms of sense organ development in *T. castaneum*, we first established a map of sensilla of the head and body segments of 1^st^ instar larvae, which were easily identifiable because of their prominent positions. The distribution of head sensilla has been described before (Schinko et al., 2008), and we therefore only define here the head sensilla relevant to this study. Each antenna has one terminal trichoid olfactory sensillum (ant_TSO; Fig. 1A, B, G, K). Long mechanosensory chaetoid sensilla on the dorsal and lateral side of the head capsule are arranged in three pairs of triplets (vertex triplet (ver_tri), gena triplet (gen_tri), maxilla escort (max_esc)) on either side of the midline, and one frontal quartet (labrum quartet (lab_qua)) (Fig. 1A, B, K; Schinko et al., 2008).

**Figure 1.**
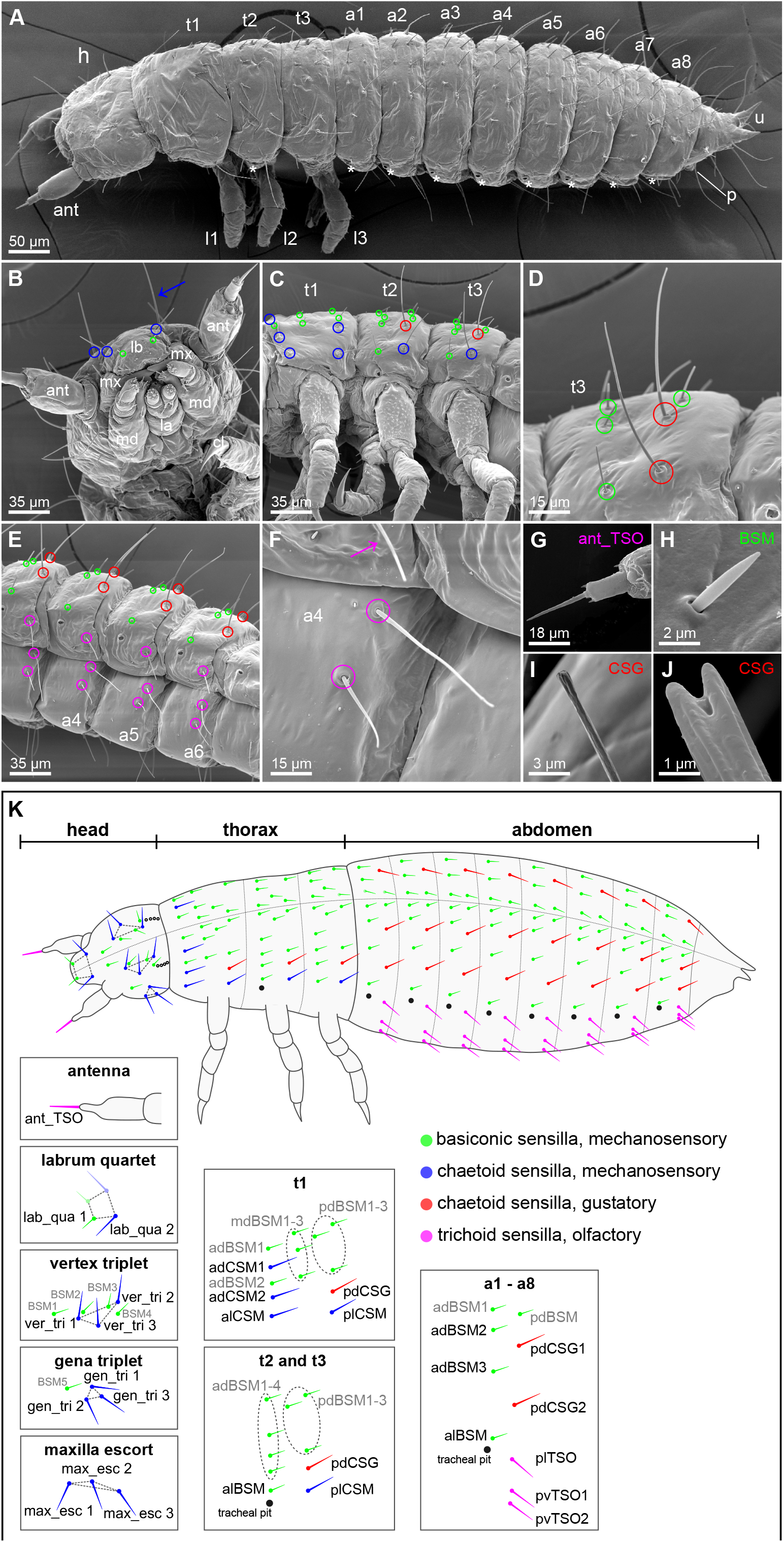
Distribution and morphology of selected external sensilla in *T. castaneum* larvae. Scanning electron micrographs (SEMs) of external sense organ morphology and distribution in the 1^st^ larval stage (A-I), and morphology of sensilla in the 2nd larval stage (J). Anterior is towards the left in A-K. Colour code sensilla: magenta, olfactory trichoid sensilla (TSOs); blue, mechanosensory chaetoid sensilla (CSMs), red, gustatory chaetoid sensilla (CSGs); green, basiconic mechanosensory sensilla (BSMs). (A) Overview of a whole larva: asterisks indicate the position of the tracheal pits; ant: antenna, h: head, t1-t3: thoracic segments 1-3, l1-l3: walking legs 1-3, a1-a8: abdominal segments 1-8, u: urogomphy, p: pygopods. (B) High magnification of the head in ventral view showing mouthparts; lb: labrum, mx: maxilla, md: mandibles, la: labium, and head sensilla which are grouped into previously described clusters lab_qua, ver_tri, gen_tri, max_esc (Schinko et al., 2008); BSMs are marked with green, and some CSMs with blue circles and arrow (see boxes in K for more details). (C and D) show thoracic segments with sensilla colour coded as in (K). (E) Ventro-lateral view of abdominal segments with magenta circles indicating TSOs. (F) High magnification of the TSOs. The S-shaped form of the TSOs is best visible in vTSO2 at this focus level. (G) High magnification of the antennal TSO (ant_TSO). (H) High magnification showing the basiconic shape of a BSM. (I) In the 1^st^ larval stage the CSGs have a bulb-shaped tip. (J) Open pore at the tip of a CSG at 2nd larval stage. (K) Schematic representation of a 1^st^ stage larvae showing the different types of external sensilla in prominent positions, dorso-lateral view. See boxes for nomenclature of sensilla, grey indicated sensilla were not analysed in RNAi screen. The dashed line indicates the dorsal midline. On t1, four CSMs are present (dCSM1-2, alCSM, plCSM) plus eight basiconic mechanosensory sensilla (adBSM1-2, mdBSM1-3, pdBSM1-3, green). In addition, one gustatory chaetoid sensillum is visible (dCSG, red). t2 and t3 show the same pattern: dCSG, plCSM and 8 BSMs (adBSM1-4, alBSM, pdBSM1-3). The abdominal segments a1-a8 display a repetitive distribution: five BSMs (adBSM1-3, alBSM, pdBSM), dCSG1-2 and three olfactory trichoid sensilla (lTSO, vTSO1-2). Sensilla abbreviations, head: ant_TSO, antennal trichoid sensillum (olfactory); lab_qua, labrum quartet; ver_tri, vertex triplet; gen_tri, gena triplet; max_esc, maxilla escort. Thorax: adBSM, anterior-dorsal basiconic sensillum (mechanosensory); alBSM, anterior-lateral basiconic sensillum (mechanosensory); alCSM, anterior-lateral chaetoid sensillum (mechanosensory); dCSG, dorsal chaetoid sensillum (gustatory); dCSM1-2, dorsal chaetoid sensillum (mechanosensory) 1-2; mdBSM1-3, median-dorsal basiconic sensilla (mechanosensory) 1-3; pdBSM1-3, posterior-dorsal basiconic sensilla (mechanosensory) 1-3; plCSM, posterior-lateral chaetoid sensillum (mechanosensory). Abdomen: adBSM1-3, anterior-dorsal basiconic sensillum (mechanosensory) 1-3; alBSM, anterior-lateral basiconic sensillum (mechanosensory); dCSG1-2, dorsal chaetoid sensillum (gustatory); lTSO, lateral trichoid sensillum (olfactory); pdBSM, posterior-dorsal basiconic sensillum (mechanosensory); vTSO1-2, ventral trichoid sensilla (olfactory) 1-2.

In the thoracic segments, the distribution of sensilla is similar in thoracic segments two and three (t2, t3; Fig. 1A, C, K). However, thoracic segment one (t1) differs from t2 and t3 as it is about twice as big (Fig. 1C) and the sensilla are arranged in three anterior-posterior rows (Fig. 1C, K, anterior, medial, posterior) rather than two. Based on the characteristics described in Table 1, we identified 5 chaetoid sensilla on t1, four of which are mechanosensory (dorsal chatoid sensilla 1 and 2 (adCSM1 and 2), anterior and posterior lateral sensilla (alCSM and plCSM)), while the remaining dorsal chaetoid sensillum is gustatory (pdCSG) (Fig. 1A, D, I, K). The two posterior sensilla pdCSG and plCSM are located in the same relative position in t2 and t3. In addition, t1 exhibits an anterior-dorsal basiconic mechanosensory sensillum (adBSM), a median transverse row of three dorsal BSMs (mdBSM1-3), and a posterior transverse row of three dorsal BSMs (pdBSM1-3; Fig. 1A, H, K). t2 and t3 show an anterior and posterior transverse row of BSMs only (adBSM1-4, alBSM, pdBSM1-3; Fig. 1A, D, K). One of the anterior BSMs, alBSM, is located in a prominent position lateral to the tracheal pit in t2 and t3 and can also be identified in all abdominal segments (Fig. 1A, C, E, K). In t1, tracheal pits are absent and the same relative position is occupied by alCSM.

Similar to t2 and t3, the sensilla are arranged in an anterior and posterior row in the abdominal segments. The anterior row consists of four BSMs (adBSM1-3 and alBSM). In the posterior row a single BSM is visible (pdBSM; Fig. 1A, E, K). In addition, each abdominal segment has five chemosensory sensilla: two chaetoid sensilla (pdCSG1 and 2) and three trichoid sensilla (plTSO, pvTSO1 and 2). pdCSG1 and 2 are located at the dorsal-posterior side of each abdominal segment and develop an open pore at the tip in the second larval stage (Fig. 1A, E, J, K). They can therefore be classified as contact chemoreceptors. One of the three trichoid sensilla is positioned at the posterior-lateral side of the abdominal segments (plTSO), at the same vertical line and posterior to the tracheal pit (Fig. 1A, E, F, K). The remaining two trichoid sensilla (pvTSO1 and 2) are located on the ventral side of the abdominal segments. The steep insertion angle and curved shape of the TSOs suggests an olfactory function.

### Expression of *Tc ASH* and *Tc ato* in the developing sense organs

In *D. melanogaster*, *ac-sc* and *ato* outline the areas where sense organs form. We therefore analysed the expression patterns of the single *T. castaneum Ac-Sc* and *ato* homologs, *Tc ASH* and *Tc ato*. In the developing peripheral nervous system, clusters of *Tc ASH* and *Tc ato* positive cells are visible in the head, thoracic and abdominal appendages as well as in domains lateral to the thoracic appendages and lateral to the developing ventral nerve cord in the abdominal segments (Fig. 2A, B; Suppl. Fig. 2). *Tc ASH* is also strongly expressed in the central nervous system (Fig. 2E; Suppl. Fig. 2A; see also Wheeler et al., 2003). In the areas of sense organ development *Tc ato* expression starts as early as NS3, while *Tc ASH* expression is not visible before NS7 (Suppl. Fig. 2B, D; stages after Biffar and Stollewerk, 2014, with additional subdivisions (Suppl. Fig. 1)). Both genes are expressed in many domains in the appendages (Fig. 2A, B; Suppl. Fig. 2C, E, F). Here, we focus on the areas from which the sense organs depicted in the scheme in Fig. 1K arise, except for the head capsule, where gene expression cannot be related to the larval sensilla.

**Figure 2.**
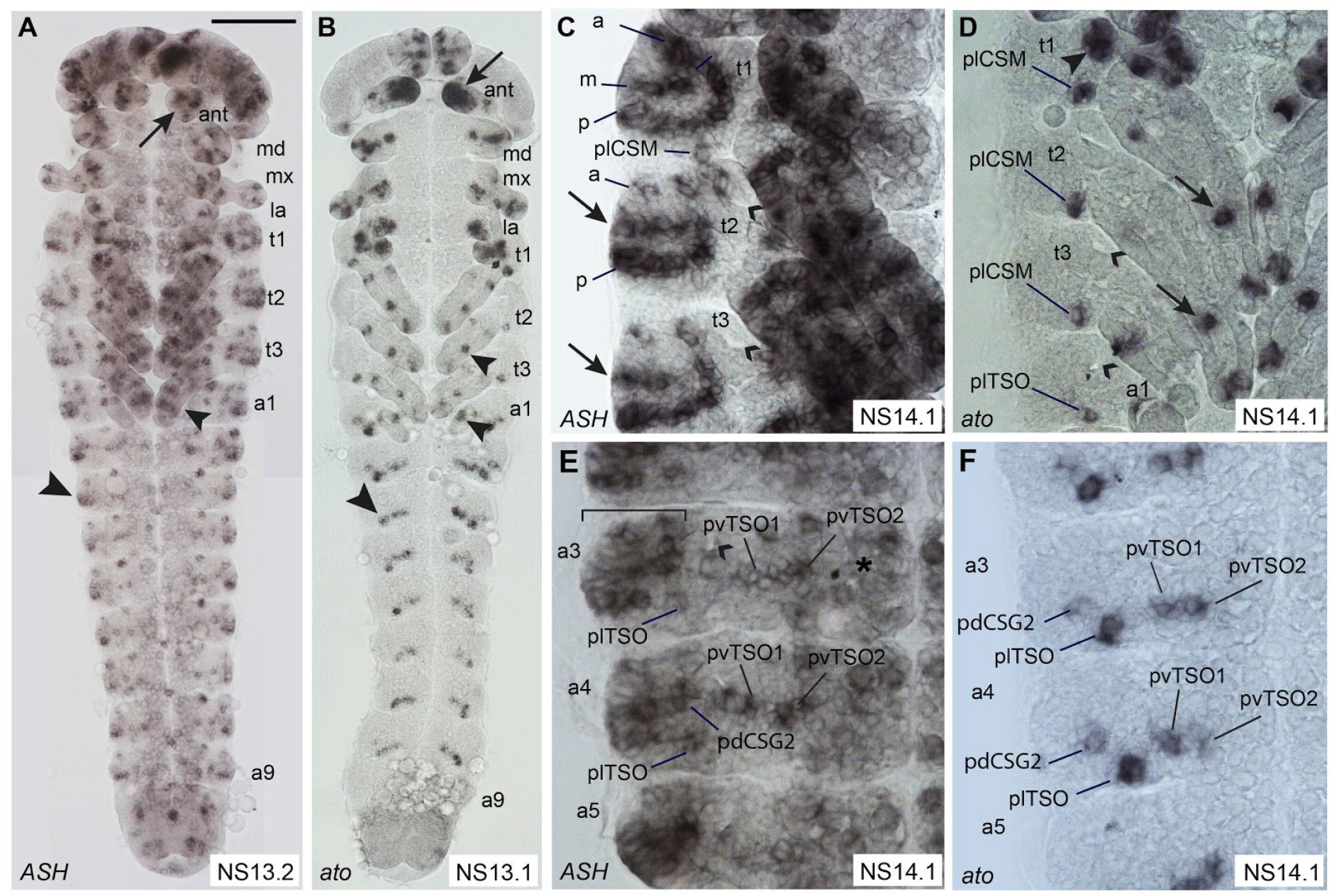
Comparison of *Tc ASH* and *Tc ato* expression patterns in the developing peripheral nervous system. Light micrographs of flat preparations stained with DIG labelled RNA probes. Anterior is towards the top. The open arrowheads point to the tracheal pits. (A-B) *Tc ASH* and *Tc ato* expression pattern in whole embryos at NS13. The arrows point towards the expression in the antennae. *Tc ato* is strongly expressed in the whole tip of the antenna, while *Tc ASH* is expressed in small groups of cells. The small arrowheads point to expression in the legs. The large arrows indicate expression in the lateral body wall. (C) In the three thoracic segments, *Tc ASH* is expressed in three dorso-ventral rows (anterior-dorsal (a), medio-dorsal (m) and posterior-dorsal (p)) dorsal to the appendages, which covers all areas from which the analysed external sensilla form. In addition, there is a medio-lateral expression domain in t2 and t3 (arrows) corresponding to an area which is devoid of external sensilla in 1^st^ stage larvae. Abundant transcripts are also visible in the legs. (D) In the thorax, *Tc ato* is expressed in a single group of cells at the posterior base of the legs, which corresponds to plCSM, in addition to a few groups in the legs (arrows). At approximately the same position, a *Tc ato* positive cluster is visible in the abdominal segments, which corresponds to plTSO. In t1, an additional large cluster of *Tc ato* positive cells is visible (arrowhead). (E) In the most lateral (dorsal) part of the abdominal segments (bracket), *Tc ASH* is expressed in many cells, covering the area from which the pdCSGs and BSMs arise. *Tc ASH* is also expressed in the ventro-lateral areas from which the three TSOs arise and in the ventral neuroectoderm (asterisk). (F) *Tc ato* is expressed in pdCSG2 and the three TSOs in the abdominal segments. For abbreviations see Fig. 1. Scale bar in A, 100 μm in A, B; 25 μm in C, D, F, G. Scale bar in E, 25 μm in E, H.

*Tc ASH* is expressed in more domains than *Tc ato*, in particular in regions lateral (dorsal) to the thoracic appendages and dorsal to the *Tc ato* domains in the abdominal appendages (bracket in Fig. 2E). In most cases, it is not possible to distinguish clusters of *Tc ASH* expressing cells belonging to individual sense organs in the dorsal domains because of their close proximity. *Tc ASH* expression in the lateral body wall starts with the areas from which alBSM and plCSM develop at NS7 to NS10 (Suppl. Fig. 2B, C). This expression persists and the pattern develops into three rows of *Tc ASH* expression (Fig. 2C: anterior, medial, posterior) dorsal to the appendages in the thoracic segments covering all positions from which larval sensilla emerge (Fig. 2C; t1: pdCSM1-2, alCSM, plCSM and all BSMs; t2 and t3: pdCSG, plCSM and all BSMs, including alBSM). The medial row in t2 and t3 (arrows in Fig. 2C) does not exhibit external sensilla in the 1^st^ larval stage. In contrast, *Tc ato* is initially only expressed in alBSM between NS7 and NS10, in addition to the appendages (Suppl. Fig. 2E, F). At NS13.1, *Tc ato* is expressed in a single cluster of cells close to the posterior base of the three thoracic appendages (Fig. 2D). This position most likely corresponds to plCSM in the larva (Fig. 1K). In addition, *Tc ato* is expressed in a large cluster of cells in t1 in the same area where the tracheal pits arise in t2 and t3 (Fig 2D). The cluster might cover the position of several sensilla (alCSM, adCSM2, adBSM2).

In the abdominal segments, the dorsal-most *Tc ASH* domain also spreads over the areas from which all described sensilla arise: adBSM1-3, alBSM, pdBSM and pdCSG1-2 (Fig. 2E). Four clusters of *Tc ASH* positive cells are furthermore visible ventral to the dorsal domains in all abdominal segments; one is located posteriorly and in approximately the same vertical line as the tracheal pit, and three are positioned in approximately the same horizontal line posteriorly to the tracheal pit. These positions correlate with the *Tc ato* positive domains that correspond to the positions of sense organs plTSO and pdCSG2, pvTSO1 and 2 in the 1^st^ larval stage (Fig. 2F). In addition, *Tc ato* is strongly expressed at the tip of the antennae, which correlates with the position of the trichoid olfactory antennal sensillum (ant_TSO) in the 1^st^ larval stage (Fig. 2B; Suppl. Fig. 2F). *Tc ASH* clusters are also visible in the antenna but not at the very tip (Fig. 2A; Suppl. Fig. 2C).

### Expression of genes conferring sense organ subtype identity in *Drosophila: Tc ct, Tc cato, Tc tap*

In order to examine sensory organ subtype specification, we next analysed the expression of *T. castaneum* homologs of genes that specify subtypes in *D. melanogaster*. These include *Tc ct*, *Tc cato* and *Tc tap*. Despite numerous attempts, we were not able to obtain in situ hybridisation data for *Tc amos*, the third member of the *T. castaneum* Atonal family and *poxn*, for which we have functional data (see below).

*Tc ct* expression starts in the maxillary and labial appendages in NS7 and becomes visible in the antennae appendages by NS10 (Suppl. Fig. 3A, B). The gene remains expressed in the labrum and whole antennae through subsequent stages (Fig. 3A, B). In addition, *Tc ct* is strongly expressed in ring-shaped domains in t2 to t3 and a1 to a8 from NS10 onward (Fig. 3E, F, I, J, open arrowheads; Suppl. Fig. 3B, C). These areas develop into the tracheal pits. In the thoracic and abdominal segments, low *Tc ct* expression becomes visible dorsal to the appendages in t1 to t3 and dorsal to the developing ventral nerve cord in the abdominal segments from stage 14.1 onward (Fig. 3E, I). At this stage, expression is also present in all appendages and the CNS (Fig. 3E, I).

**Figure 3.**
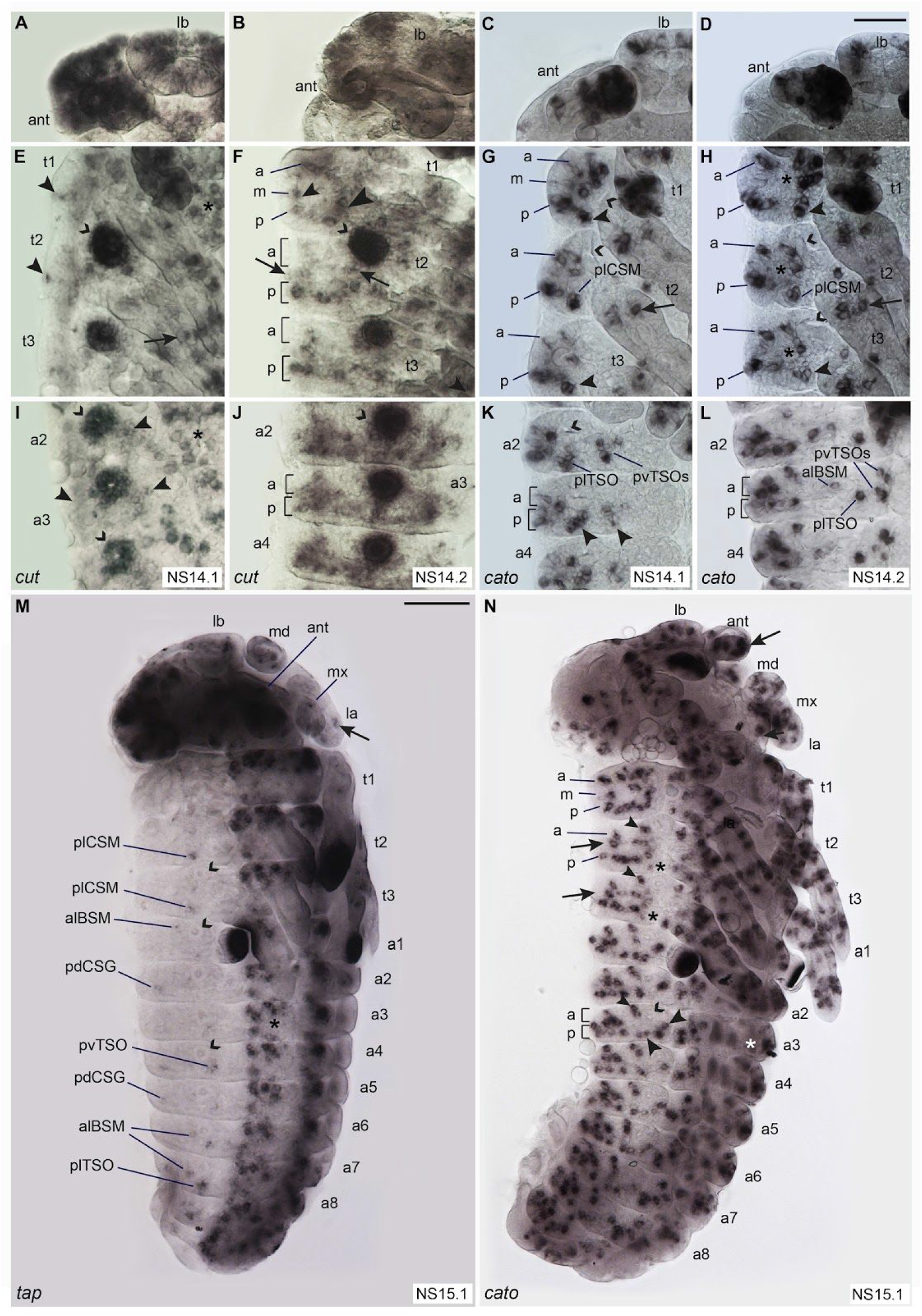
Comparison of the expression patterns of sense organ subtype specific genes. Light micrographs of flat preparations stained with DIG labelled RNA probe of *Tc ct*, *Tc cato* and *Tc tap*, respectively. The open arrowheads point to the tracheal pits. (A-D) Both *Tc ct* and *Tc cato* are expressed in the labrum and the antennae, although *Tc ct* covers larger areas than *Tc cato*. (E) At NS14.1, *Tc ct* is expressed in a few cells lateral to the appendages in the thorax. *Tc ct* is also expressed in the ventral neuroectoderm (asterisk). (F) At NS14.2, *Tc ct* positive cells are arranged in a ring-like structure in t1 covering the area from which the anterior-dorsal (a), and posterior-dorsal (p) rows of larval setae arise; however, in the medial row (m) staining is only visible in two clusters, one in the most dorsal position (small arrowhead) and one close to the appendages (large arrowhead). In t2 and t3, *Tc ct* positive cells are aligned in a posterior-dorsal row and at least two clusters of *Tc ct* expressing cells are visible in the medio-dorsal row (arrows). The area covering the anterior row of larval setae shows only scattered, slightly stained cells. (G) At NS14.1, *Tc cato* is expressed in clusters of cells which can be roughly allocated to a, m and p rows in the thoracic segments. The plSCM cluster is clearly visible due to its prominent position posterior to the tracheal pits and close to the appendages (arrowheads). (H) At NS14.2, *Tc cato* is expressed in two rows (a, p) and a medial cluster of cells (asterisks) in the thoracic segments. The plCSM clusters (arrowheads) are identifiable in t1 to t3. (I) At NS14.1, *Tc ct* is expressed dorsally, below, and ventrally to the developing tracheal pits in the abdominal segments and in the ventral neuroectoderm (asterisk). (J) At NS14.2, *Tc ct* is expressed in the whole area from which the posterior row of larval setae arises in the abdominal segments, while the anterior row is only partially covered. (K) At NS14.1, *Tc cato* is expressed in clusters of cells in the a and p rows. The plTSO and pvTSO1 and 2 clusters can be distinguished due to their prominent positions posterior and ventral to the tracheal pits, respectively (arrowheads). (L) At NS14.2 plTSO, pvTSO1 and 2 and alBSM are identifiable in addition to the remaining a and p *Tc cato* positive clusters. (M) At NS15.1, *Tc tap* is expressed in the thoracic plCSMs, thoracic and abdominal alBSMs as well as in some of the abdominal pdCSGs. In addition, the gene is expressed in the head and thoracic appendages (arrows) and the tip of the abdominal appendages. (N) At NS15.1, the arrangement of *Tc cato* positive cells in the a, m and p rows is more pronounced due to the dorso-ventral extension and anterior-posterior narrowing of the segments. The expression in the thoracic plCSM clusters has decreased (asterisks); however, *Tc cato* positive cell clusters corresponding to the positions of alBSM (small arrowheads) as well as plTSO and pvTSO1 and 2 (large arrowheads) are still identifiable. In addition, *Tc cato* remains expressed in the ventral neuroectoderm and the head and thoracic appendages are covered in clusters of *Tc cato* positive cells. A large domain of *Tc cato* is visible at the tip of the antennae and the pleuropods. For abbreviations see Fig. 1. Scale bar in D 25 μm in A-L; scale bar in M, 100 μm in M, N.

In the peripheral nervous system, the fully developed *Tc ct* expression pattern can be observed in NS14.2 (Fig. 3B, F, J). Similar to *Tc ASH*, *Tc ct* expression forms an incomplete ring-shape in t1, although covering a smaller region (Fig. 3F). However, in the *Tc ASH* positive area corresponding to the medio-dorsal row of setae in the larva (mdBSM1 to 3; Fig. 2C), only one *Tc ct* cluster is visible in the dorsal-most position that might correlate to expression in mdBSM1 (Fig. 3F, small arrowhead). There is no expression in the remaining part of the row, except close to the base of the first thoracic leg (Fig. 3F, large arrowhead). This area cannot be directly correlated with the position of external larval setae. In t2 and t3, *Tc ct* expressing cells are arranged in a posterior row lateral to the appendages but appear scattered towards anterior (Fig. 3F). Similar to t1, two clusters of *Tc ct* positive cells are visible in the medial area in both t2 and t3, however, these cannot be correlated with external larval setae (Fig. 3F, arrows). In the abdominal segments, the fully developed *Tc ct* expression pattern covers the complete posterior row of larval setae, including the lateral and ventral TSOs medial to the tracheal pits (Fig. 3J). The anterior and posterior expression domains form a continuous area, which takes on a triangular shape towards anterior and thus does not cover the positions of all larval setae in the anterior row (Fig. 3J).

In the peripheral nervous system, *Tc cato* is first expressed in the antennae and mandibles in stage NS 7 embryos (Suppl. Fig. 3D). In subsequent stages additional *Tc cato* expression sites form in the antennal, maxillary, labial and thoracic appendages (Fig. 3C, G; Suppl. Fig. 3F). From stage NS10 onward, clusters of *Tc cato* expressing cells become visible lateral to the appendages in the thorax and lateral to the ventral neuroectoderm in the abdominal segments (Suppl. Fig. 3F). With the appearance of additional expression domains, the clusters become arranged into rows (Fig. 3G, H, K, L, N). The developing plCSM, alBSM, plTSO and pvTSO1 and 2 sensilla are clearly visible as separate clusters, while the remaining clusters merge into each other and are not identifiable relative to the position of the larval sensilla (Fig. 3G, H, K, L, N). The arrangement of *Tc cato* positive cells into rows is more pronounced in stage NS15.1, possibly due to the segments having narrowed along the anterior posterior axis and extended along the dorso-ventral axis as part of germband retraction and dorsal closure (Fig. 3N). The posterior rows of *Tc cato* expression in the thoracic and abdominal segments seem to cover all larval sensilla positions, except for plCSM, where the previous expression has almost ceased. Three clusters in the medial row of t1 might correspond to the medio-dorsal BSMs1-3 (Fig. 3N). Similar to *Tc ASH* expression, a medial row of expression is also visible in t2 and t3, which cannot be correlated to external larval sensilla (Fig. 3N). In t2, t3 and the abdominal segments, there is a gap of *Tc cato* expression in the medial part of the anterior rows. In abdominal segments, however, several *Tc cato* positive cell groups cluster in the dorsal part of the anterior row, which might rearrange to cover the gap during further dorso-ventral extension of the germband (Fig. 3N). Furthermore, many *Tc cato* clusters are visible in all appendages and there is a strong expression domain at the tip of the antennae (Fig. 3N).

*Tc tap* is strongly expressed in the CNS from NS11 onward (Suppl. Fig. 3E); however, expression is comparably low in the peripheral nervous system. It does not start before NS15.1 (Fig. 3M) and begins to decrease after about 6 hours in NS15.4 (Suppl. Fig. 3G). *Tc tap* is expressed in clusters of cells in the head appendages and in a few cells in the lateral body wall of the thorax and the abdomen (Fig. 3M). *Tc tap* positive cells are identifiable in the plCSMs, alBSMs, plTSOs and pvTSOs as well as in one of the two abdominal pdCSGs (Fig. 3M; Suppl. Fig. 3G).

### Expression patterns of *Tc asense*, *Tc prospero* and *Tc snail*

In *D. melanogaster ase*, *pros* and *sna* are so-called panneural genes, which are expressed in all sense organs after SOP formation. *Tc ase* shows a strong and prolonged expression in the CNS (Fig. 4A; Wheeler et al., 2003; Biffar and Stollewerk, 2014); however, the expression is limited to a transient expression in a few cells and clusters in the PNS (Fig. 4A). Expression in the PNS starts at NS9 in small domains on the appendages and a few cells on each side of the lateral body wall (Suppl. Fig. 4A).

**Figure 4.**
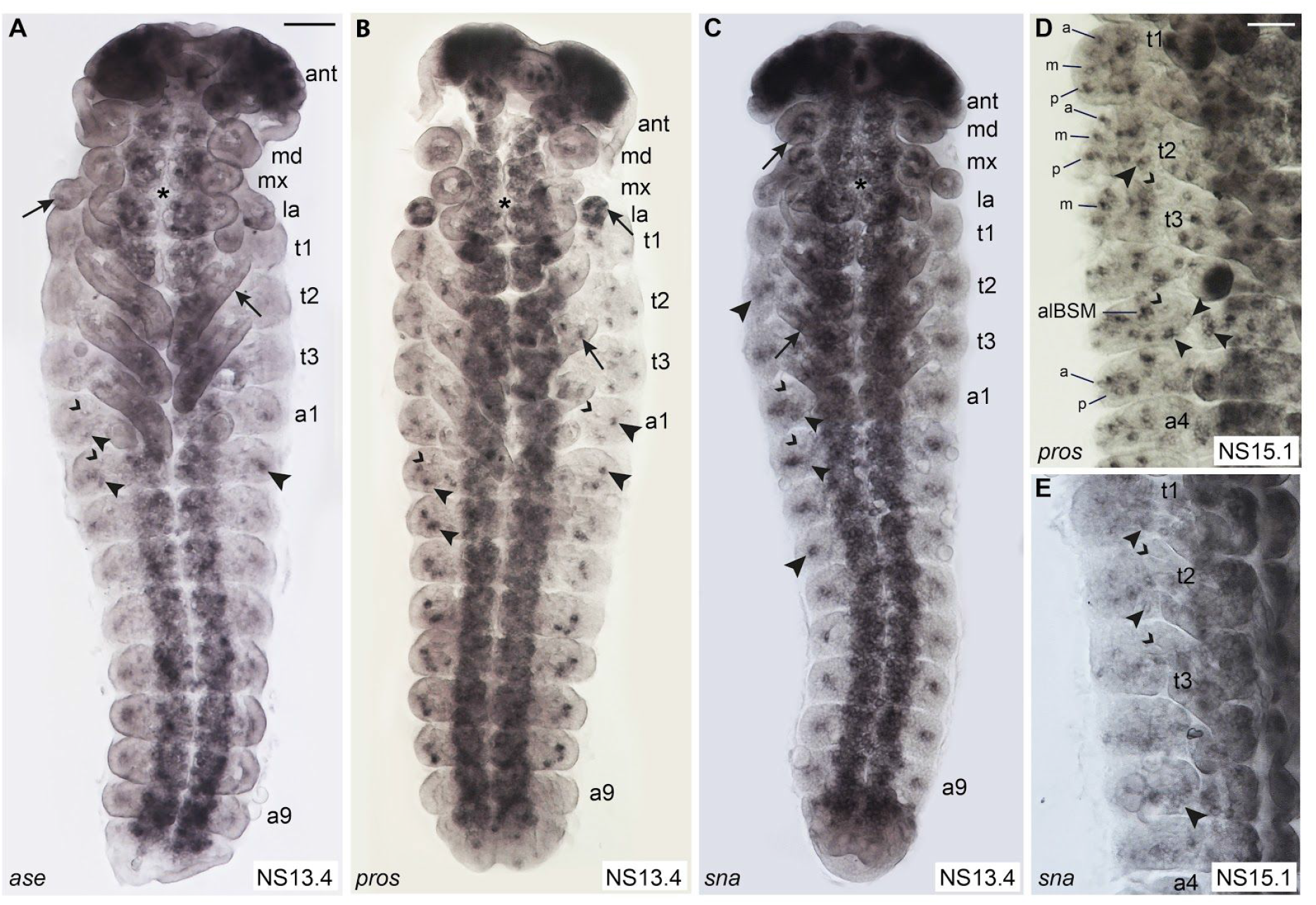
Comparison of the expression patterns of panneural genes. Light micrographs of flat preparations stained with DIG labelled RNA probe of *Tc ase*, *Tc pros* and *Tc sna*, respectively. The open arrowheads point to the tracheal pits. (A) *Tc ase* is strongly expressed in the developing brain, the antennae and the ventral neuroectoderm (asterisk). Scattered cells express *Tc ase* in the remaining appendages (arrows) and the lateral body wall. The stained cells directly posterior to the tracheal pits might belong to the developing plTSOs (arrowheads). (B) *Tc pros* is expressed in the same area below the tracheal pits (small arrowheads) indicating that it is co-expressed with *Tc ase* in the plTSOs. *Tc pros* expression dorsal to the tracheal pits most likely corresponds to the alBSMs (large arrowheads). *Tc pros* is also strongly expressed in the developing brain, in clusters of cells in all appendages (arrows) as well as in the ventral neuroectoderm (asterisk). (C) Similarly, *Tc sna* is expressed in the brain, clusters of cells in the appendages (arrows) and ventral neuroectoderm (asterisk). In addition, *Tc sna* is expressed in large clusters in the lateral body wall of each segment (large arrowheads). The expression extending below the tracheal pits might correspond to the developing plTSOs (small arrowheads). (D) At NS15.1, groups and single cells express *Tc pros* in the lateral body wall, which seem to cover all areas of external sensilla formation. Additionally, the medial row that does not give rise to external sensilla expresses *Tc pros*. Due to their prominent positions relative to the tracheal pits, the alBSM clusters, the plCSM clusters (large arrowhead) and the three TSO clusters (small arrowheads; plTSO, pvTSO1 and 2) are easily identifiable. (E) *Tc sna* shows a transient expression pattern in groups and single cells, some of which cover the areas where sensilla appear next to landmarks, such as the plCSMs (small arrowheads) in the thoracic segments and the plTSOs in the abdominal segments (large arrowhead). For abbreviations see Fig. 1. Scale bar in A, 50 μm in A-C; scale bar in D, 50 μm in D, E.

At NS11 additional clusters appear in the abdominal segments (Suppl. Fig. 4B). One of the clusters can be identified as plTSO after formation of the tracheal pits in the abdominal segments at NS13 (Fig. 4A). The expression domain located dorsal and posterior to the tracheal pits cannot be assigned to specific sensilla (Fig. 4A). *Tc ase* expression decreases thereafter and is not detectable any more in most areas of the PNS by NS15 (Suppl. Fig. 4F).

*Tc pros* expression is first visible in the head appendages at NS7 (Suppl. Fig. 4C). At NS10, a single cluster of *Tc pros* positive cells appears on each side of the lateral body wall (Suppl. Fig. 4G). By NS13.4 additional clusters are present, three of which can be attributed to the developing plCSMs (thorax) and plTSOs (abdomen) and the alBSMs (Fig. 4B). At NS15.1, *Tc pros* positive clusters seem to cover all areas of external sensilla formation (Fig. 4D). In addition, *Tc pros* is expressed in the medial area in t2 and t3 that does not give rise to external sensilla (Fig. 4D). Due to their prominent positions around the tracheal pits, alBSM, plCSM, plTSO and pvTSO1 and 2 can be clearly identified (Fig. 4D).

*Tc sna* expression starts at NS7 in the mandibles and by NS9 all appendages show *Tc sna* expression domains (Suppl. Fig. 4D). Similar to *Tc pros*, bilateral *Tc snail* positive clusters appear in the lateral body wall. They appear first in the thoracic segments at NS7 and have extended to A9 by NS9 (Suppl. Fig. 4D, E). One additional *Tc sna* expression domain is visible at NS13, which can be allocated to the abdominal plTSO due to its position posterior to the tracheal pits (Fig. 4C). During the subsequent stages, *Tc sna* shows a transient expression pattern in groups and single cells, some of which cover the areas where sensilla appear next to landmarks, such as the plCSMs in the thoracic segments (Fig. 4E). *Tc sna* expression decreases earlier than that of *Tc pros* in the PNS, although both genes continue to be strongly expressed in the CNS (Fig. 4D, E).

### *Tc ASH* and *Tc ato* have different roles in sense organ development

The expression of *Tc ASH* and *Tc ato* in SOPs raises the possibility that these genes specify sense organ subtype like in *D. melanogaster*. In order to test this hypothesis, we disrupted *Tc ASH* and *Tc ato* function via parental RNAi and examined sense organs in affected larvae. We focused our analysis on a subset of clearly identifiable sensilla as described for the 1^st^ larval stage in Fig 1.

When examining control larvae (from parents injected with water or buffer alone), 98 to 100% of these sensilla were present (Fig. 5A; Suppl Table 3; Fig. 6A). The observed variation is due to the absence of sensilla at specific positions along the anterior-posterior axis. In the head and thorax, 99.1% of the analysed sensilla are present at all positions (2,484/2,507), while in the abdominal segments overall 2.87% (199/6912) of sensilla are missing. The abdominal TSOs (4.32%) show the highest variability followed by the CSGs (2.60%). In order to elucidate the significance of the RNAi phenotypes, we recorded and analysed the affected sensilla separately for the head, thorax and abdomen (Fig. 5; Suppl. Fig. 6). For Fig. 5, we put the numbers of affected sensilla together for each sensilla category (TSOs, CSMs, BSMs, CSGs) across all larvae analysed.

**Figure 5.**
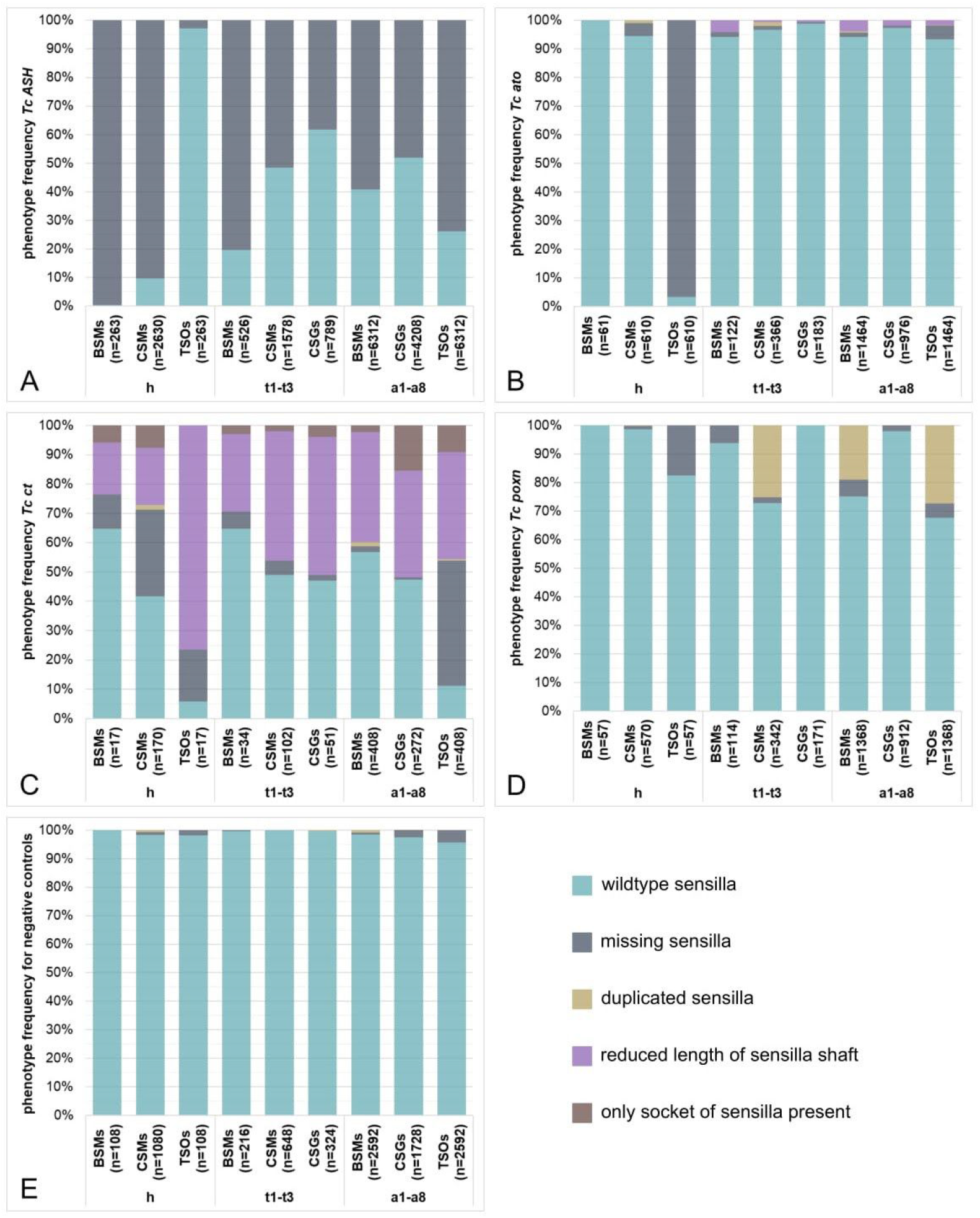
Quantification of the RNAi phenotypes of external larval sensilla. The bars represent the percentages of phenotypes identified for the different ESO subtypes (BSMs, CSMs, CSGs and TSOs) on the head (h) and the thoracic (t1-t3) and abdominal segments (a1-a8) of *Tc ASH* (A), *Tc ato* (B), *Tc ct* (C), *Tc poxn* RNAi (D) and the negative control cuticles (E), respectively. Sensilla that were not affected are categorised as ‘wildtype’ (turquoise). Sensilla showing a phenotype are divided into the four categories ‘missing sensilla’ (grey), ‘duplicated sensilla’ (beige), ‘reduced length of sensilla shaft’ (purple) and ‘only socket of sensilla present’ (brown), if applicable. (A) We analysed 387 specimens for both non-overlapping dsRNA fragments (NOF1 and 2) of *Tc ASH* larvae in total, of which 263 showed a specific phenotype (sensilla missing, grey). (B) We analysed 119 specimens of *Tc ato* RNAi (NOF1 and NOF2 collectively), of which 61 showed a phenotype. *Tc ato* RNAi cuticles were missing the ant_TSOs (96.72%, see 3rd bar, grey), and also observed a small percentage of duplicated sensilla (beige) and sensilla with reduced shaft length (purple). (C) We were able to analyse only 26 specimens in total for both NOFs of *Tc ct* (n=17 showed a phenotype). Sensilla of *Tc ct* cuticles could be grouped into the four different categories of phenotypes. The most abundant phenotype for all ESOs types was identified as ‘sensilla with reduced shaft length’ (purple). (D) We performed pRNAi in *T. castaneum* pupae to examine the function of *Tc poxn*. We analysed 111 specimens in total. 51.35% of the analysed specimens showed a phenotype which were identified as duplicated sensilla (CSMs on thorax, BSMs and TSOs on abdomen) (beige). See Suppl. Table 2 for summary of RNAi injection results.

**Figure 6.**
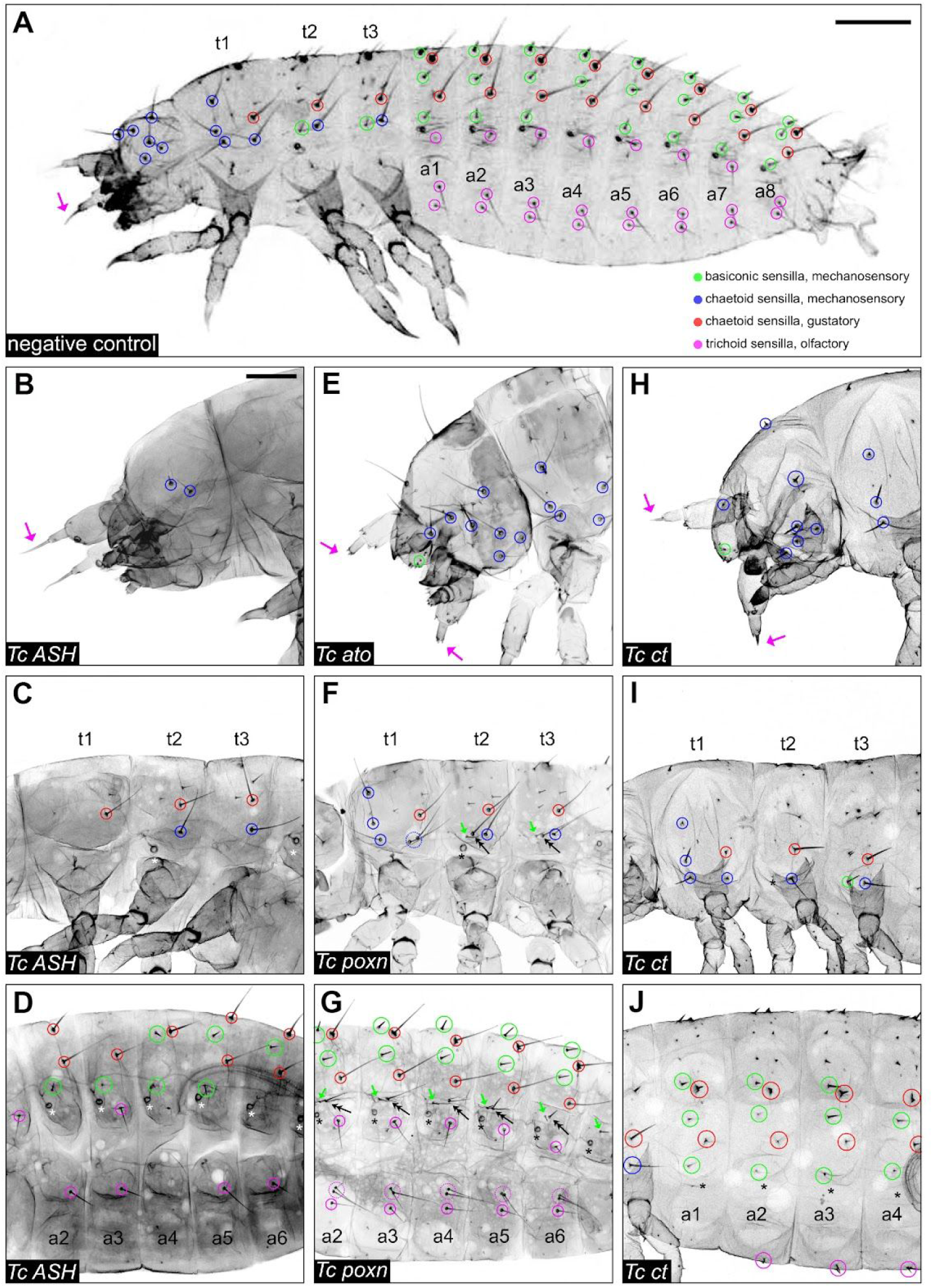
RNAi phenotypes of cuticles. Laser-scanning confocal images of L1 cuticles; anterior is to the right. The gene targeted by RNAi is indicated in the lower left corner. (A) The colour coded sensilla analysed in this study are indicated in a negative control cuticle for comparison. (B) On the head of the *Tc ASH* RNAi cuticle only few CSGs (blue circles) and the ant_TSO (magenta arrow) are present. (C) On the thoracic segments of *Tc ASH* RNAi cuticles only pdCSG on t1, plCSMs and pdCSGs on t2 and t3 are present. The remaining CSMs and the two BSMs are missing. (D) On the abdominal segments of the *Tc ASH* RNAi cuticle all three types of sensilla (BSMs, CSGs, and TSOs) are affected. Sensilla in different positions are missing, e.g. on a1, adBSM1 and 2, plTSO, and pvTSO2 are missing. (E) The *Tc ato* RNAi cuticle of the head shows missing antennal flagella, ant_TSOs (magenta arrows). (F and G) The *Tc poxn* RNAi cuticle shows duplications of specific sensilla (plCSM, alBSM and pvTSO1) on thorax and abdomen. (F) On the thorax the plCSMs (dotted blue circle) are duplicated, and an additional sensilla is found between alBSM and plCSM on t2 and t3, which has the morphological characteristics of plCSMs (black arrows). (G) On the abdomen (a2-a8) pvTSO1s are duplicated in *Tc poxn* RNAi cuticles (dotted magenta circles). An additional sensilla is found posterior to the alBSMs in a1 to a6 (black arrows). The additional sensilla exhibit a longer shaft compared to the wildtype BSMs. (H) On the head of *Tc ct* RNAi cuticles, the ant_TSOs (magenta arrows) and CSMs and BSMs have shorter shafts (green and blue circles). ver_tri1 and 2 are missing. (I) On t1, adCSM1, plCSM and pdCSG develop a socket only. On t2, the alBSM is missing. (J) The shaft of the CSGs (red circles) and alBSMs (green circles) are shorter or only developed as a socket. plTSOs are missing, and pvTSOs have a shorter shaft (magenta circles). Asterisks in (I and J) indicate missing tracheal pits. Scale bar in A, 100 μm; scale bar in B, 50 μm.

In *Tc ASH* RNAi larvae, two types of morphologically distinct mechanosensory sensilla (basiconic and chaetoid) are missing from the head: 90.38% of the CSMs and 99.62% of the BSMs (Fig. 5A; Fig. 6B, C, D). In addition, 2.73% of the olfactory antennal trichoid sensillum ant_TSO are missing, however, this is only slightly above the proportion of defects seen in control larvae (1.84%; Fig. 5D; Suppl. Table 3). A similar pattern is seen on the thorax, where BSMs are most affected (80.42% missing) and to a lesser extent the CSMs (51.52% missing) (Fig. 6C). In addition, 38.15% of the gustatory chaetoid sensilla, the CSGs, are absent on the thorax (Fig. 5A).

Although the head TSO pattern is not significantly changed, the TSOs are the most affected sensilla in the abdominal segments. 73.80% of the TSOs (plTSOs, pvTSOs 1 and 2) are missing in the abdomen, followed by 59.28% of the BSMs and 48.05% of the CSGs (Fig. 5A; Fig. 6D). In the most severe *Tc ASH* RNAi phenotypes all sensilla are missing (Suppl. Fig. 6D, F, J). In addition to the sensilla phenotypes, the pretarsal segments of the legs, the urogomphi, and the mandibles appear rounded (Suppl. Fig. 6B, H, L compared to Suppl. Fig. A, G, K). The ‘rounded pretarsal segment’ phenotype has also been documented in the iBeetle screen and in another insect (Schmitt-Engel et al., 2015; Finet et al., 2018). Taken together, our results clearly show that *Tc ASH* is required for the generation of all subtypes of ESOs analysed, except for the antennal TSOs.

In contrast to *Tc ASH*, *Tc ato* has only a minor role in the formation of larval ESOs. The antennal TSOs are the only ESOs that are frequently missing (96.72%) in *Tc ato* RNAi (Fig. 5B, dark blue bar; Fig. 6E). All other ESOs are present on the head, and on the thorax and abdomen, BSMs, CSMs, CSGs and TSOs are only missing to a small percentage, which is in the range of the variations seen in the controls (0.55-4.92%; Fig. 5B). The differentiation of all types of sensilla is also affected at a small rate in the thorax and abdomen: the sensilla are duplicated or have shorter shafts compared to wildtype (0.1-1.37%, beige and 0.55-4.10%, purple in Fig. 5B).

### Functional analysis of the ‘subtype’ identity genes

Next we analysed the role of the *T. castaneum* homologs of genes that specify subtype identity in *D. melanogaster*, namely *Tc ct*, *Tc poxn, Tc amos*, *Tc cato* and *Tc tap.* Parental *Tc ct* RNAi resulted in sterile females and we therefore performed embryonic RNAi to examine gene function. In *Tc ct* RNAi larvae, all types of ESOs are affected (Fig. 5C; Fig. 6H, I, J). However, compared to the *Tc ASH* phenotype, most of the sensilla are present in *Tc ct* RNAi larvae but they show differentiation defects. We categorised these defects into ‘duplicated sensilla’, ‘reduced length of sensilla shaft’, ‘only socket of sensilla present’, ‘sensilla missing’, and ‘wildtype’ (Fig. 5C). The TSOs are most strongly affected in *Tc ct* RNAi. On the head, the ant_TSOs are affected in 94.12% of the cases, while 89.22% of the abdominal TSOs that show abnormal shape (Fig. 5C, turquoise). The predominant phenotype for all TSOs is a reduction in sensilla length (76.47% of ant_TSOs and 36.76% of abdominal TSOs; Fig. 6H, J, magenta arrows and circles).

The CSMs are the second most affected ESO subtype in *Tc ct* RNAi larvae: 58.24% show a phenotype in the head and 50.98% in the thorax (Fig. 6H, I; blue circles). There is a difference in the distribution of the sensilla phenotypes between head and thorax. In more than half of the affected CSM sensilla positions in the head, sensilla are absent (29.41%), while reduced length of sensilla shaft is the predominant phenotype in the thoracic CSMs (44.12%, Fig. 5C). Furthermore, 52.57-52.94% of the thoracic and abdominal CSGs are affected (Fig. 6I). The most prominent phenotype is again the reduced length of the sensilla shaft (36.40-47.06%). However, the abdominal CSGs show the highest percentage of the ‘only socket of sensilla present’ phenotype (15.44%) compared to all other sensilla types (Fig. 5C, brown bars). The BSMs are about equally affected across all body parts (35.29% in the head and thorax, 43.14% in the abdomen; Fig. 5C; Fig. 6I, J). Again there are slight differences in the distribution of the sensilla phenotypes. In the head, about one third of the BSMs are missing in the affected positions, while differentiation defects are predominant in the affected BSMs of the remaining body parts (Fig. 5C). In addition, tracheal pits are absent in 96.15% of the *Tc ct* RNAi larvae, which is in line with the prominent circular expression of the gene in the areas where the tracheal pits develop (Fig. 3E, F, I, J, open arrowheads; Suppl. Fig. 3B, C, Fig. 6I, J; black asterisks indicating missing tracheal pits). Taken together, *Tc ct* is required to various degrees for the specification development of all ESO subtypes.

In contrast to the *Tc ct* RNAi phenotype, only a small subset of sensilla are affected in *Tc poxn* RNAi larvae. In t1 to t3, the mechanosensory sensillum plCSM is duplicated (50.88%) and in the abdominal segments the mechanosensory alBSM (55.92%) and the olfactory pvTSO1 (81.80%) show the same duplication phenotype (beige bars in Fig. 5D; double arrowhead in Fig. 6F and G, magenta dotted circles in Fig. 6G).

We also attempted to analyse a gene that is regulated by Poxn in *D. melanogaster*, *target of pox neuro* (*tap*); however, *Tc tap* RNAi larvae were not analysable because of gross morphological defects such as the absence of the abdomen or deformation of head and thorax. These results are in line with those of the iBeetle screen (Schmitt-Engel et al. 2015). Similarly, we also attempted to functionally analyse two additional members of the Atonal family, *Tc amos* and *Tc cato*. However, similar to *Tc tap* RNAi, severe structural defects made it impossible to analyse the sensilla pattern in *Tc cato* RNAi larvae.

## Discussion

In this study we analysed several genes in *T. castaneum*, homologs of which determine or change the fate of sense organ precursors in *D. melanogaster.* Here we discuss their divergent roles in the two insect species and the implications on the evolutionary scenario of sense organ diversification in arthropods.

### Proneural gene expression and sense organ subtype identity are not directly linked in *T. castaneum*

In *D. melanogaster*, there are essentially 5 bHLH transcription factors (*ac, sc, ase, ato, amos*) that combined account for the formation of all sense organs (Hartenstein, 2005). A striking discovery of our study is that in *T. castaneum* a single transcription factor, *Tc ASH*, is required for the formation of all morphological and functional classes of ESOs analysed. This includes olfactory trichoid sensilla (TSOs), gustatory chaetoid sensilla (CSGs) and two types of mechanosensory sensilla, chaetoid (CSMs) and basiconic (BSMs). In contrast, in *D. melanogaster*, Achaete (Ac) and Scute (Sc) determine the identity of external mechanosensory (chaetoid, trichoid, campaniform) and gustatory sensilla, excluding olfactory sense organs (Campuzano et al., 1985; Cubas et al., 1992; Reddy et al., 1997).

Olfactory sense organs are determined by Ato and Amos in *D. melanogaster*, which are required for different subsets of these sensilla. In *ato* mutants the olfactory coeloconic sensilla on the antennae and the basiconic sensilla on the maxillary palps are absent (Gupta and Rodrigues, 1997), while Amos is required for the remaining olfactory sensilla on the antennae, the olfactory basiconic and trichoid sensilla (zur Lage et al., 2003). In contrast, in *T. castaneum*, *Tc ato* is only required for the formation of one of the analysed olfactory sense organs, ant_TSOs (96.72% ant_TSOs absent in *Tc ato* RNAi), while the abdominal TSOs appear unaffected. *T. castaneum* head appendages bear small numbers of coeloconic, basiconic and additional trichoid sensilla (Ryan and Behan 1973, Behan and Ryan 1978), and future studies may reveal a role for *Tc ato* in these olfactory sense organs. We also observed a reduction in length in all sensilla in *Tc ato* RNAi, including in those that do not appear to express *Tc ato*. While this could be due to off-target/toxic effects of the dsRNA, we favour the hypothesis that *Tc ato* could function during sensilla morphogenesis at a late stage of development that was not captured in our analysis.

In addition to the olfactory sense organs, *D. melanogaster ato* is required for the formation of the chordotonal organs, which are internal (mechanosensory) stretch receptors (Jarman et al., 1993a, 1995). Larval chordotonal organs develop during embryogenesis in an invariant pattern in the lateral body wall (Dambly-Chaudiere and Ghysen, 1986). Although there is no information about larval chordotonal organs in beetles, it can be assumed that they are distributed across the body wall similar to *D. melanogaster*. Surprisingly, *Tc ato* expression is limited to areas that give rise to external sensilla suggesting that it does not play a role in larval chordotonal organ formation in *T. castaneum*. However, it is tempting to speculate that the expression of *Tc ASH* in the medial row of t2 and t3, which does not give rise to external sensilla (Fig. 2C, arrows), might correspond to the formation of internal sense organs.

Our data show that similar to *D. melanogaster ac*, *sc* and *ato* (Campuzano et al., 1985; Cubas et al., 1992; Reddy et al., 1997), *Tc ASH* and *Tc ato* are the first genes to be expressed in the SOPs and are thus at the top of the gene cascade. Sense organs requiring these gene products for their formation are absent in the corresponding functional studies. However, in contrast to *D. melanogaster*, the *T. castaneum* proneural genes do not directly link the formation of sense organ precursors with the acquisition of subtype identity since *Tc ASH* is required in all sense organ types and *Tc ato* specifies a single (or few) olfactory sense organ(s) rather than all sense organs of this type. Furthermore, *Tc ato* is expressed in subsets of chaetoid and basiconic mechanosensory (alBSM, alCSM, plCSM) as well as gustatory chaetoid sensilla (pdCSG2), although the functional relevance is not clear (see below). This raises the question of at which level and how subtype identity is determined in *T. castaneum*.

### The regulation of subtype identity genes has diverged in *T. castaneum*

In *D. melanogaster,* the determination of sense organ subtype identity seems to be a decision between two (or three) alternative developmental pathways. Misexpression and loss-of-function studies show that Ac-Sc promote external mechanosensory versus internal mechanosensory (chordotonal organ) fate, while Ato supports formation of olfactory versus gustatory/mechanosensory sensilla externally, in addition to internal mechanosensory organ versus external mechanosensory organ fate (Rodriguez et al., 1990; Brand et al., 1993; Dominguez and Campuzano, 1993; Hinz et al., 1994; Jarman et al., 1993a; Jarman and Ahmed, 1998). *Ct*, a sense organ subtype specific gene acting downstream of Ac-Sc, seems to play a central role in determining the alternative fates. Misexpression of *D. melanogaster ct* in chordotonal organs of the lateral body wall transforms these organs into external mechanosensory organs and external mechanosensory organs develop into chordotonal organs in *ct* mutants (Blochinger et al., 1991; Bodmer et al., 1987; Meritt, 1997). Vice versa, misexpression of *ato* in external mechanosensory organs of the notum or wing result in a transformation to chordotonal organs (Jarman and Ahmed, 1998). These data show that both Ato and Ct establish sense organ subtype identity and can individually switch SOP fate in *D. melanogaster*. In contrast, Sc is not able to transform sense organ fate when misexpressed (Jarman and Ahmed, 1998).

It was shown in the chordotonal organs of *D. melanogaster* that Ato exerts its subtype determining role by suppressing the activation of *ct,* a gene necessary for external sense organ fate (Jarman and Ahmed, 1998). This allows switching to the alternative developmental pathway of chordotonal organ formation. *D. melanogaster ct* is also not expressed in olfactory sense organs (Jhaveri et al., 2000). Thus *ato* and *ct* expression are mutually exclusive (except for Johnson’s organ, see below). In contrast in *T. castaneum, Tc ct* and *Tc ato* are co-expressed, indicating that *Tc ato* does not repress *Tc ct*. A collaboration of *Tc ato* and *Tc ct* in sense organ development is supported by the fact that in *Tc ct* RNAi the antennal olfactory sense organ ant_TSO, which requires *Tc ato*, is affected to a similar percentage as in *Tc ato* RNAi (*Tc ct* RNAi, 94%; *Tc ato* RNAi, 97%). However, the predominant phenotype in *Tc ct* RNAi is a reduction in sensilla length, rather than the loss of the sensilla, indicating that *Tc ct* is involved in the differentiation of ant_TSO.

We also observed a strong *Tc ct* RNAi phenotype in the remaining TSOs that coexpress *Tc ato*, *Tc ct* and *Tc ASH*, the abdominal plTSO and pvTSO1 and 2. *Tc ct* is expressed later than *Tc ato* and *Tc ASH* suggesting that the gene might be activated by one of these proneural genes. Although not substantially affected in *Tc ato* RNAi, these olfactory sensilla show a high percentage of differentiation defects in *Tc ct* RNAi (89%) and are frequently absent in *Tc ASH* RNAi (74%). *Tc ct* is expressed in all types of analysed sensilla and the predominant phenotype in *Tc ct* RNAi are differentiation defects, such as shorter sensilla and missing sensilla shafts (‘sockets only’, Fig. 5), rather than transformation into other sensilla subtypes. These data suggest that we have to strike off another gene from the list of potential subtype identity genes in *T. castaneum*: *Tc ct* does not confer subtype identity, rather, the gene product might play a role in sensilla morphogenesis in all sense organs.

We tested four additional genes for potential roles in subtype identity: *poxn*, *tap*, *amos*, *cato*. Poxn is required in the polyinnervated ESOs of larval and adult *D. melanogaster* (Dambly-Chaudiere et al., 1992). Although the function of polyinnervated sensilla is not known in all cases, the ones that have been studied are of the gustatory type, such as the recurved gustatory bristles on the anterior wing margin, the labellum and the legs (Jiang et al., 2015). In *poxn* mutants wing discs, the recurved bristles at the anterior wing margin are transformed into monoinnervated mechanosensory bristles (Awasaki and Kimura, 1997). Similarly in *poxn* mutant larvae, the three types of polyinnervated sensilla (two ‘kölbchen’ per thoracic hemi-segment; one papilla and hair per abdominal hemi-segment) are transformed into monoinnervated mechanosensory organs (Jiang et al., 2015). In contrast, we did not observe a transformation of sensilla in *T. castaneum* larvae. If we assume that in *T. castaneum*, *Tc tap* is activated by *Tc poxn* and expressed in the same polyinnervated sensilla as is the case in *D. melanogaster* (Gautier et al., 1997), Tc Poxn is only involved in the development of one to four sensilla per hemi-segment: plCSM in t1, plCSM and alBSM in t2 and t3 and alBSM, plTSO, pvTSO, and one of the two dCSG in the abdominal hemi-segments. In *Tc poxn* RNAi, three types of these sensilla are duplicated: plCSM, alBSM and pvTSO1. The duplication might be due to a duplication of the SOPs or within the SOP lineage. The latter is supported by a detailed study of the sensilla lineages in *D. melanogaster poxn* mutant larvae showing that Poxn is required in the progeny of the SOPs and regulates the number and types of cells produced by each secondary precursor (Jing et al., 2015). Functional studies of *Tc tap* could have contributed to our understanding of the subtype identity mechanisms in *poxn* positive sense organs but unfortunately, the *Tc tap* RNAi larvae were not analysable. However, even in *D. melanogaster*, the role of *tap* remains elusive, as only the overall PNS neuron and glia arrangement was analysed in *tap* mutant embryos and the pattern was found unchanged (Yuan et al., 2016).

Interestingly, in t2 and t3 the duplicated plCSM sensilla in *Tc poxn* RNAi larvae are closer in position to the alBSM sensilla. Such a shift in position has also been observed for transformed sensilla in the anterior wing margin rows of *D. melanogaster poxn* mutant wings but the underlying cause is unknown (Awasaki and Kamura, 1997).

Taken together, we found no obvious role of Tc Poxn in subtype identity since we neither observed a transformation of one sensilla type into another in *T. castaneum* larvae nor loss or differentiation defects in specific morphological or functional sensilla types. However, we cannot rule out that the affected sensilla share characteristics that were not considered here, such as a similar innervation pattern. This needs to be resolved by future morphological and cell lineage studies. We attempted to analyse the function of the Atonal Family genes *Tc amos* and *Tc cato*. However, severe overall morphological defects prevented us from studying the sensilla pattern. Based on our gene expression analysis *Tc cato* does not show sense organ subtype specific expression: it is expressed in all morphological and functional types of sensilla. Expression starts after formation of the SOPs, slightly earlier than that of *Tc ct*. However, in *D. melanogaster*, *cato* expression is also not restricted to a specific sense organ type: *cato* expression is initially only visible in the Ato and Amos dependent sense organs, but later it appears also in the external mechanosensory sensilla after *ac-sc* are switched off and after the start of *ct* expression (Goulding et al., 2000a). *Cato* mutants show duplications of sensory neurons (zur Lage and Jarman, 2010). The severe morphological defects following *Tc cato* RNAi must be due to functions during earlier stages of development than those we studied.

### Evolutionary scenario

Our data show that in contrast to *D. melanogaster*, the concept that the identity of categories of sense organs can be assigned to the function of single or combinations of transcription factors cannot be applied in *T. castaneum*. Regardless of how we categorize the sense organs - by morphology, by function or both - there is no transcription factor code that can be aligned with a single category. This raises the question of what can be considered as the ancestral pattern in insects.

Based on data from other arthropod groups and within insects, we can assume that Achaete-Scute and Atonal Family members played a role in sense organ development in the last common ancestor of insects (e.g. Klann and Stollewerk 2017; Pioro and Stollewerk 2006; Tadesse et al., 2011; Stollewerk and Seyfarth, 2008). Data on neurogenin-related genes (*tap* in *D. melanogaster*), *poxn* and *ct* in sense organ development are not available in other arthropod groups. However, we show here for the first time that these genes are involved in sense organ development in an insect other than *D. melanogaster* suggesting that they might have belonged to the sense organ toolkit in the last common ancestor of insects.

In order to understand how the determination of sense organ subtype identity has diverged in the two insects, we need to consider the roles of the sensory genes in the process of sense organ specification. The specification of sense organ identity can be subdivided into three subsequent events: formation and subtype specification of SOPs, specification of accessory (e.g bristles, sockets) and neural cell types, and differentiation of sense organ cells. In both insects, the sensory genes are active at different time points and thus can be assigned different roles in the process of subtype identity development based on their expression and mutant phenotypes. Both in *T. castaneum* and in *D. melanogaster*, *Tc ASH* and *ac-sc,* respectively, are expressed at the top of the gene cascade during formation of the SOPs; however, in *T. castaneum, Tc ASH* does not function as a subtype specific gene in contrast to *D. melanogaster*, rather, it is required in all functional and morphological subtypes of sense organs. Based on the presented data, we suggest the following evolutionary scenario. In the ancestral state, *ASH* is the predominant proneural gene for sense organ development. ASH endowed epidermal cells with the potential to develop into external sense organs *without* simultaneously specifying subtype identity. Consequently, the sense organs developed along a default pathway relying on ‘general’ cell fate specification and differentiation genes that are expressed in all sense organs (e.g. *sna*, *pros*, *Notch*, *numb*). The default state might have been overwritten by the recruitment of additional proneural genes. This assumption is supported by the *D. melanogaster amos* mutant phenotype. *Amos* is required for subsets of olfactory sense organs on the antennae. In *amos* mutants, ectopic mechanosensory bristles appear in place of olfactory sensilla (zur Lage et al., 2003). This phenotype is accompanied by a derepression of *ac-sc* expression in the SOPs of the ectopic bristles, indicating that Amos controls subtype identity by repressing *ac-sc* in the wildtype condition. Similarly, *D. melanogaster* Ato promotes chordotonal organ formation via repressing the downstream target of Ac-Sc *ct* (Jarman and Ahmed, 1998). The hypothesis that additional proneural genes were recruited to increase sense organ diversity is further supported by the co-expression of *Tc ASH* and *Tc ato* in several sensilla; however, how these two genes collaborate in sense organ specification needs further investigation. A widespread co-expression of ASH and ato is also seen in crustacean sense organ development suggesting that this evolutionary strategy is used across pancrustaceans (see Klann and Stollewerk, 2017 for further discussion).

Additional genes that were recruited to sensory development, might have appeared late in development (e.g. *tap*) and were initially only required for regulating neuronal differentiation such as axonal projections and the structure of dendrites. Other genes were recruited to the earlier process of accessory (socket, shaft, sheath cells) and neuronal cell type specification (neurons, glia) to regulate the developmental potential and divisions of the secondary precursors (e.g. *cato*, *poxn*). The addition of sensory genes to the existing set resulted in variations in cell type numbers and morphology that were used as an evolutionary tool for sense organ diversification.

In the next step, changes in the temporal expression of the add-on genes changed their role and importance in sense organ development. There are several examples in *D. melanogaster* that support this evolutionary scenario. *D. malanogster tap*, for example, is expressed at a late stage of gustatory sense organ formation in the gustatory neuron and its precursor, indicating a role in neuronal differentiation. However, during development of the adult antennal olfactory sensilla, *tap* is expressed at the time of SOP formation (Ledent et al., 1998). Similarly, *cato* shows variations in its temporal expression in different sense organs. In a single chordotonal organ (lch5), the gene is required for the specification of the SOP, while it is involved in sense organ cell type specification in all other chordotonal organs (zur Lage and Jarman, 2010). In ESOs, *cato* is expressed later than in the chordotonal organs and up until terminal differentiation (Blochinger et a., 1990). Furthermore, Ct, which is expressed in ESO SOPs, has a late role in dendritic arborization in a subtype of multidendritic (MD) neurons (Ebacher et al., 2007). Moreover, although not expressed in any of the other chordotonal organs, Ct plays a role in the differentiation of the scolopidia in the fly’s auditory Johnson’s organ, which are the individual units of chordotonal organs. Similar to *ato* mutants, *D. melanogaster ct* mutants are deaf, however, in *ato* mutants scolopidia are not formed at all (Ebacher et al., 2007). Thus, there is one sense organ in *D. melanogaster*, which requires both *ato* and *ct* for its development.

The latter example points out another consequence of the temporal expression shift: the escape from constraining upstream regulations. In all chordotonal organs, except Johnson’s organ, Ato executes its subtype identity by suppressing *ct* expression (Jarman and Ahmed, 1998). However, due to its late expression, *ct* can be engaged as a differentiation gene in Johnson’s organ. Similarly, Cato can control neuron numbers in Ac-Sc dependent sense organs after these genes have been switched off (zur Lage and Jarman, 2010).

Taken together, the examples from *D. melanogaster* support an evolutionary scenario whereby sensory genes are recruited to the development of individual or subsets of sense organs, which do not necessarily fall into specific morphological or functional classes. Changes in the temporal expression can move the genes up in the hierarchy so that they can control all aspects of a specific sense organ subtype, as is the case for *ct* and *poxn* in *D. melanogaster*, for example. The *T. castaneum* data presented here also fit this evolutionary scenario. *Tc ct* has been recruited to a late step of development in all sense organs, regulating the differentiation of shaft cells, while Tc Poxn is only required in a subset of sensilla possibly controlling cell type numbers. *Tc cato* is expressed earlier than *Tc cut* and might therefore be involved in sensory organ cell type specification, similar to *Tc poxn* but in all sense organs. None of these genes are expressed during formation of the SOPs and therefore do not have control over the whole specification process, i.e. sense organ subtype identity.

To conclude, the evolutionary scenario presented here suggests that sense organ diversity has evolved from a default state through subsequent recruitment of sensory genes to the different sense organ specification processes. A role in subtype identity has evolved as a secondary effect of the function of these genes in individual or subsets of sense organs. It is obvious that in this scenario prepatterning genes (which we have not discussed here but see e.g.: Rozowski and Akam, 2002, Eksi et al., 2018) would have had a major influence on the evolution of the positional identity of sense organs. Future comparative analysis will show how patterning mechanisms can be fit into the evolution of sense organ diversity.

## Material and Methods

### Animal husbandry and gene sequences

*T. castaneum* beetles were reared as previously described (Brown et al. 2009). The San Bernadino (SB) wildtype strain was used for all experiments. Sequences were amplified from cDNA (synthesised from extracted RNA from different developmental stages with SuperScript III First-Strand Synthesis System (Invitrogen) and cloned into pGEM®-T Easy Vector (Promega) using standard cloning procedures.

### mRNA probe synthesis and in situ hybridisation

Antisense mRNA probes for genes in this study were synthesised from their cloned sequences (as described above). The in vitro transcription using the T7 RNA polymerase and the DIG RNA labelling Mix (both Roche) was performed following the supplier’s protocol. Colorimetric whole mount in situ hybridisation (NBT/BCIP) was performed as previously described using anti-DIG antibody conjugated with Alkaline-Phosphatase (Roche) (Schinko et al. 2009).

### RNA interference and statistical analyses

Parental RNAi (pRNAi) (*Tc ASH*, *Tc ato*, *Tc poxn*, *Tc cato*, *Tc tap*, *Tc amos*) and embryonic RNAi (eRNAi) (*Tc ct*) were performed to infer gene functions. The gene sequences were obtained from the iBeetle-Base (Dönitz et al. 2015, Dönitz et al. 2018). Double stranded RNA (dsRNA) for all genes was ordered from Eupheria Biotech (Dresden, Germany). For each gene, two non-overlapping fragments (NOF1 and NOF2) were injected (where NOF1 is the same as the iB-RNA fragments used in the iBeetle screen (Schmitt-Engel et al. 2015; see Suppl. Tab. 1 and Suppl. Fig. 5 for list of iB-RNA numbers and location). For pRNAi, female pupae were injected with 1μl/μg dsRNA as previously described (Bucher et al. 2002, Bucher and Klingler 2004, Posnien et al. 2009 and ‘The Beetle Book’, http://wwwuser.gwdg.de/∼gbucher1/tribolium-castaneum-beetle-book1.pdf). Embryos were injected with 3μl/μg dsRNA as previously described (Benton et al. 2018). The protocol was adjusted slightly. Embryo preparation, mounting, and injection was performed as described (Benton et al., 2018). However, after injection, the coverslips with the embryos were placed upside down onto the oxygen permeable membrane with 3 stacked coverslips used as bridges, and the intervening space was filled with halocarbon oil. This petri dish set-up was inverted and placed on a layer of 1% agarose gel to maintain humidity. The embryos were kept in this condition at 32°C until the embryos hatched. For pRNAi and eRNAi experiments, negative controls were either injected with H_2_O or injection buffer (1.4 mM NaCl, 0.07 mM Na2HPO4, 0.03 mM KH2PO4, 4 mM KCl, pH 6.8).

First instar larvae (L1) of pRNAi and eRNAi experiments were used for cuticle analysis and prepared as described before (Wohlfrom et al. 2006). L1 cuticle preparations were analysed for wildtype larvae (wt), ‘phenotype’, ‘non-specific’ (i.e. broken cuticles, larvae still in egg membrane and hence sensilla not analysable, etc.), and ‘empty eggs’ (results are summarized in Suppl. Table 2). The larvae showing a phenotype were screened for the sensilla in prominent positions described in Fig. 1. These sensilla were counted in each larva on one side (preferably right side) and categorized, where applicable, as ‘wildtype’, ‘missing sensilla’, ‘duplicated sensilla’, ‘reduced length of sensilla shaft’, ‘only socket of sensilla present’. The recorded data for each gene are summarized for dsRNA NOF1 and NOF2, and further categorized into sensilla subtypes (TSOs, BSMs, CSMs and CSGs) and body parts (head, thorax, abdomen). An overview of the recorded data can be found in Suppl. Tables 3-7). Microsoft Excel was used to document and process data for statistical analysis.

### SEM sample preparation

*T. castaneum* 1^st^ instar larvae (24h egg lays were incubated at 32°C for 3 days) were collected and washed with PBS. The washed larvae were fixed in 1:1 heptane and 3% glutaraldehyde at room temperature for 1h. The fixative was removed, and the larvae were dehydrated with a series of acetone (70%, 80%, 2x 90%, 2x 100%). HMDS was used to dry the larvae (larvae were incubated in HMDS, then the solution was removed, followed by air drying larvae in a block dish overnight under the fume hood). The dried larvae were mounted on aluminium stubs with sticky tape and sputter coated with gold (Agar auto sputter coater model 108A). A FEI Quanta 3DFEG or a FEI Inspect F electron microscope was used for imaging L1 larvae visualizing sensilla morphology.

### Microscopy and image processing

Following in situ hybridization (colorimetric and fluorescence), the yolk of *T. castaneum* embryos was removed. The embryos were then mounted flat onto microscope slides using glycerol. Colorimetric stained embryos (NBT/BCIP staining reaction), as well as cleared cuticles of *T. castaneum* larvae were imaged, or screened using an inverse Leica microscope (DM IL, Wetzlar Germany) and corresponding LAS software (version 2.8.1). Image acquisition of fluorescent labelled staining, as well as L1 cuticles were performed with a Leica SP5 confocal microscope (Wetzlar, Germany) and corresponding LAS X software. Z-stacks of L1 confocal images were processed using the Z-projection tool of Fiji (Schindelin et al., 2012). Different graphic design programs (Adobe photoshop CS2, Adobe illustrator and Inkscape version 0.92) were used for image processing, assembly, and schematic illustrations preparations.

## Acknowledgements

We thank Elke Küser, Claudia Hinners and Gregor Bucher for providing SB *Tribolium castaneum* beetles and for permitting the use of the injection equipment at the Georg-August University in Göttingen and Vicente Araullo-Peters from the NanoVison Centre at QMUL for support taking SEM images.

