## Supplementary Material for "Sense organ formation and identity are controlled by divergent mechanisms in insects"

### Supplementary Figures

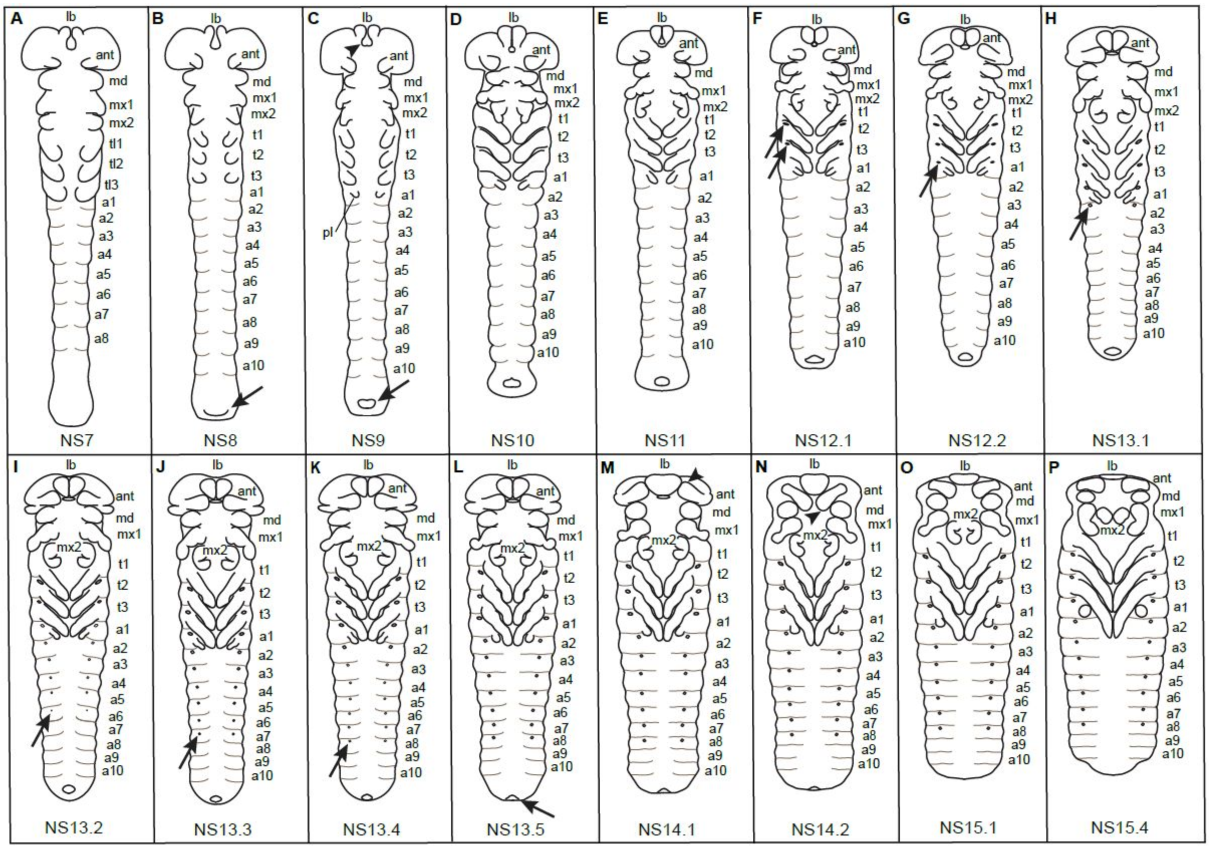


**Supplementary Figure 1.** ***Tribolium castaneum*** **staging system.** The figure shows schematic drawings based on light microscopic images of flat preparations. The schemes reflect the relative size of the embryos. In *T. castaneum,* the phases of embryonic development are usually subdivided into hours of development after egg laying (AEL) at 32 °C (Wheeler et al., 2003; Savard et al., 2006; Benton et al., 2013). *T. castaneum* development takes approximately 70h at 32°C (Brown et al., 2009). However, in order to compare gene expression patterns, in particular in RNAi treated animals where development might be delayed, a morphological staging system was required spanning the whole time of sense organ development. Biffar and Stollewerk (2014) described all stages covering central nervous system development (NS1 to NS15 corresponding to 9h to 52h AEL at 32°C). We adopted this staging system but introduced finer subdivisions of the later stages (NS12-15). This was required to capture and compare the successive development of sense organs in different preparations. The stages relevant to sense organ development (NS7-15) are depicted here. We only describe features that are easily identifiable and were used to distinguish between the stages. The hours given in brackets correspond to the time after egg laying (AEL) at 32°C, when the embryos were collected (A) NS7 (18h): limb buds are visible, the maxillae show a triangular shape and the labrum appears as a paired lobe. Three thoracic segments and eight abdominal segments are visible. (B) NS8 (20h): All 10 abdominal segments and the proctodeum (arrow) have formed. The thoracic legs have further lengthened to partially cover the posterior part of the segment. (C) NS9 (22h): The embryo has reached its greatest length. Labrum and stomodeum form a triangular shape (arrowhead). The pleuropodia, which are the appendages of the first abdominal segment are visible. The proctodeum has a round shape (arrow). (D) NS10 (24h): The legs have lengthened further so that they almost touch at the ventral midline. Indentations on the head appendages (mandible, maxilla, labium) indicate the formation of segments (podomeres). The two lobes of the labrum are aligned along their proximo-distal length. (E) NS11 (26h): Some leg pairs touch each other at the ventral midline. The gnathal appendages have elongated further and appear more differentiated. The labium has a hook-like shape. (F) NS12.1 (28h): The embryo appears shorter due to the condensation of the segments along the anterior-posterior axis and broadening along the dorso-ventral axis. Tracheal pits have appeared on the $2^{nd}$ and $3^{rd}$ thoracic segments (arrows). (G) NS12.2 (30h): The antennal tips point upwards. Tracheal pits are also visible on the $1^{st}$ abdominal segment (arrow). (H) NS13.1 (32h): The legs have grown further so that they extend into the next posterior segment. Tracheal pits are present on the $2^{nd}$ abdominal segment (arrow). (I) NS13.2 (34h): Tracheal pits appear up to the $6^{th}$ abdominal segment (arrow). (J) NS13.3 (36h): Tracheal pits are visible in the $7^{th}$ abdominal segment (arrow). (K) NS13.4 (38h): Tracheal pits are present in the $8^{th}$ abdominal segment. (L) NS13.5 (40h): The proctodeum (arrow) has moved to a more dorsal position. (M) NS14.1 (42h): The distal parts of the antennae point upwards towards anterior (arrowhead). (N) NS14.2 (44h): The distal parts of the antennae point towards posterior (arrowhead). The legs have extended to reach over two subsequent segments posteriorly. (O) NS15.1 (46h): The antennae point towards the ventral midline. The anterior-posterior length of the labrum is reduced. The embryo shortens further and extends along the dorso-ventral axis. (P) NS15.4: The embryo shortens further. The maxillae assume a hook-like shape and the labia have moved anteriorly between the maxillae. Abbreviations from anterior to posterior: lb, labrum; ant, antennal segment; md, mandibular segment; mx, maxillary segment; la, labial segment; t1-3, thoracic segment 1-3; a1-8, abdominal segment 1-8.

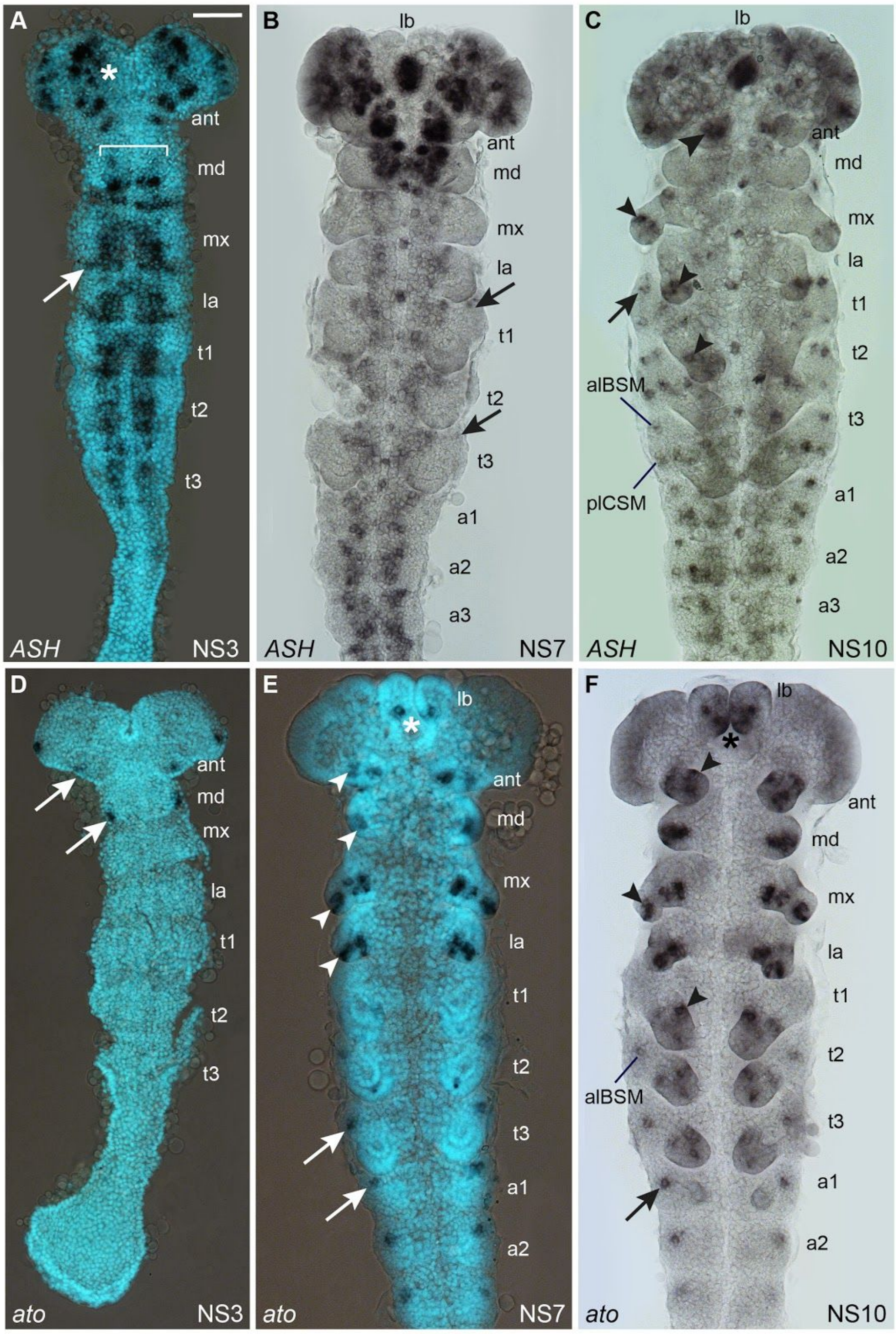


**Supplementary Figure 2. Comparison of *Tc ASH* and *Tc atonal* expression patterns in the developing peripheral nervous system.** Light (B, C, F) and fluorescence (A, D, E) micrographs of flat preparations stained with DIG labelled RNA probes and SYBR Green (light blue; A, D, E). Anterior is towards the top. (A) At NS3, *Tc ASH* is strongly expressed in the CNS (asterisk) and ventral neuroectoderm (bracket). In the thoracic segments, *Tc ASH* expression extends laterally forming a small stripe (arrow). However, these stripes disappear before clusters of cells emerge where sense organs form. (B) At NS7, *Tc ASH* expression is visible in a few cells in the lateral body wall (arrows). In addition, the gene is strongly expressed in the developing brain and ventral nerve cord. (C) At NS10, *Tc ASH* expression has increased in the peripheral nervous system and is visible in single and groups of cells in the lateral body wall (arrow). Based on the relative expression with regard to the thoracic legs and the anterior-posterior position in the segment, the expression domains correspond to the developing sense organs alBSM (anterior-lateral basiconic seta, mechanosensory) and plCSM (posterior-lateral chaetoid seta, mechanosensory). Additional *Tc ASH* domains are also visible in the antennal (large arrowhead), maxillary, labial and thoracic appendages (small arrowheads). (D) *Tc ato* is first expressed at stage NS3 in the antennal and mandibular segment (arrows). The bilateral domains in the head lobe most likely correspond to the developing optic lobes. (E) At NS7, additional *Tc ato* expression can be seen in all head appendages (arrowheads) and the labrum (asterisk). Expression starts in the developing sense organs of the lateral body wall (arrows). (F) At NS10 the same expression domains are visible in the elongated head appendages (arrowheads), the labrum (asterisk) and the lateral body wall (arrows). The expression in the lateral body wall most likely corresponds to alBSM. Additional *Tc ato* domains are visible in the thoracic appendages. Abbreviations see Suppl. Fig. 1. Scale bar: 50 µm.

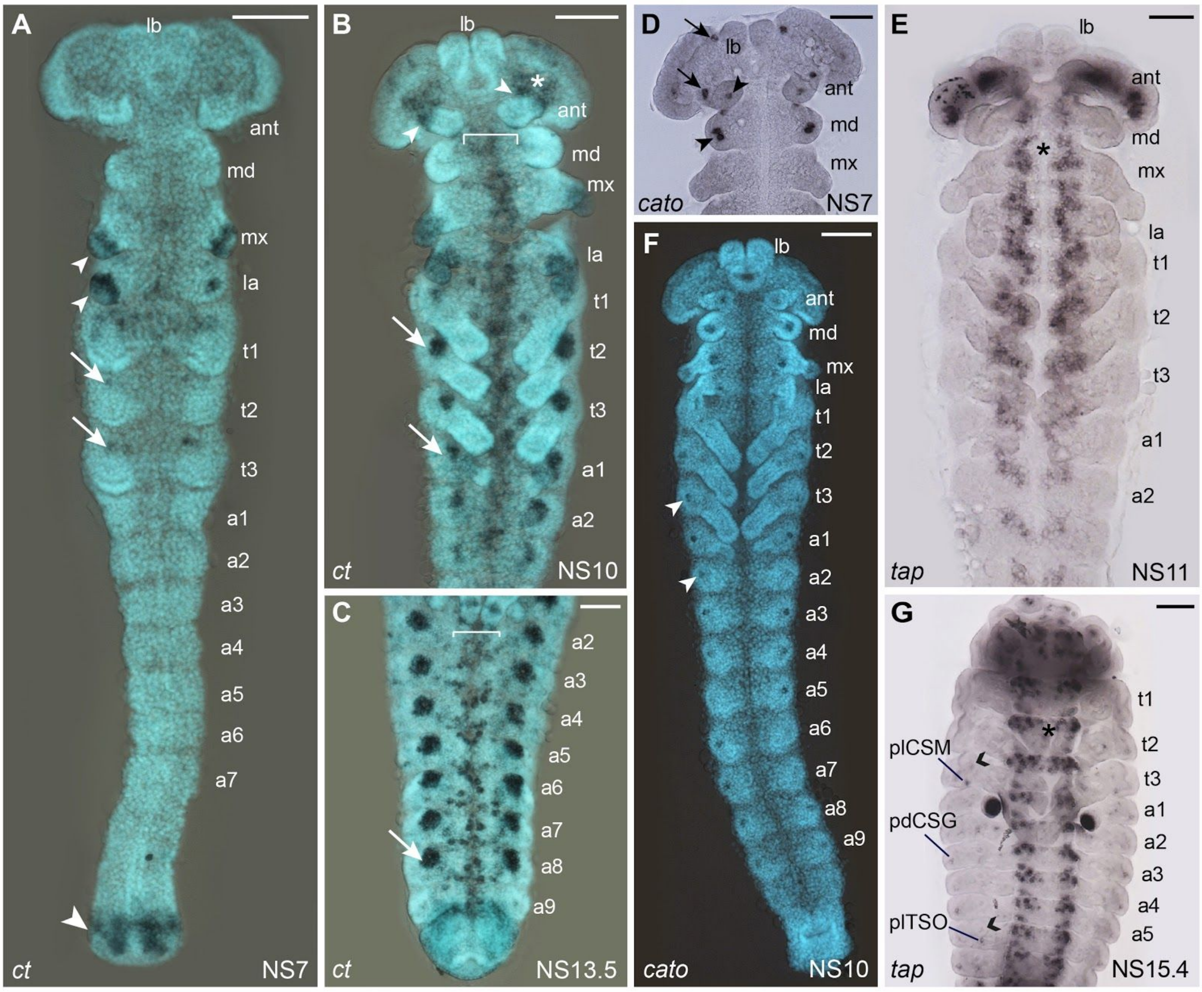


**Supplementary Figure 3. Expression patterns of the sense organ subtype specific genes *Tc ct, Tc cato* and *Tc tap.*** Fluorescence micrographs of flat preparations stained with DIG labelled RNA probes (dark blue) and SYBR Green (light blue; A-C, F) and light micrographs stained with DIG labelled RNA probes (D, E, G). Anterior is towards the top. (A) *Tc ct* expression is first visible at NS7 at the tips of the maxillary and labial appendages (small arrowheads) and around the proctodeum (large arrowhead). Small expression domains are visible in the areas where the tracheal pits form (arrows). (B) At NS10, *Tc ct* is expressed at the base of the developing antennae (arrowheads) and the whole tips of the maxillary and labial appendages. In addition, *Tc ct* expression is visible in the brain (asterisks) and the developing ventral nerve cord (bracket). *Tc ct* is also strongly expressed in the developing tracheal pits of the thoracic and first three abdominal segments (arrows). (C) At NS13.5, *Tc ct* expression is visible in the ventral neuroectoderm (bracket). The expression around the tracheal pits has extended to the 8th abdominal segment (arrow). (D) At NS7, *Tc cato* is expressed in the antennal and mandibular appendages (arrowheads). Additional expression is visible both at the base of the labrum and the antennae (arrows). (E) *Tc tap* expression starts in the CNS (asterisk) at NS11. (F) *Tc cato* is first expressed in the lateral body wall in single clusters (arrowheads) at NS10. (G) At NS15.4, *Tc tap* expression has decreased and is only detectable in a few cells corresponding to the positions of plCSM, one of the two abdominal pdCSGs and plTSO. The open arrowheads point to the tracheal pits. Abbreviations see Suppl. Fig. 1. Scale bars in A, B, G, F 100 µm; C-E, 50 µm.

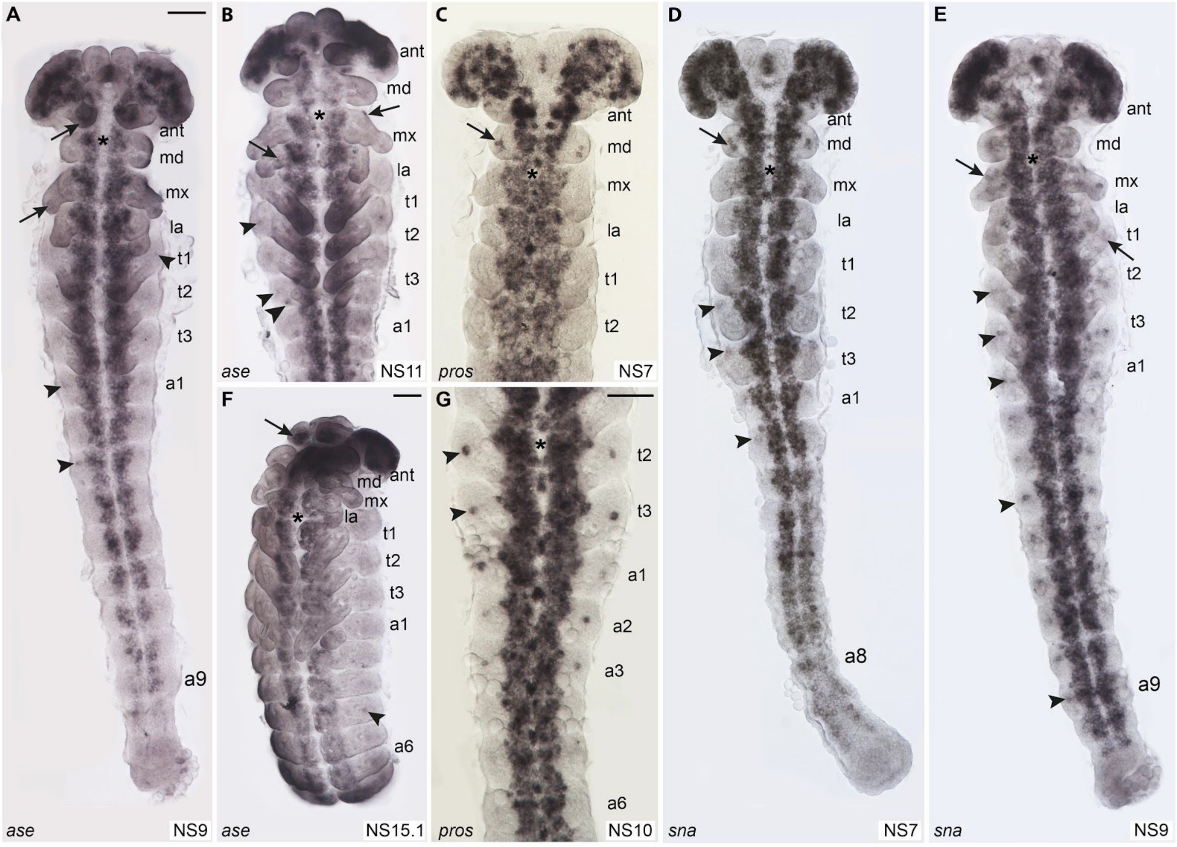


**Supplementary Figure 4. Expression patterns of the panneural genes *Tc asense*, *Tc prospero* and *Tc snail*.** Light micrographs of flat preparations stained with DIG labelled RNA probes of *Tc ase*, *Tc pros* and *Tc sna*, respectively. Anterior is towards the top. The asterisks indicate the expression of the genes in the CNS. (A) At NS9, *Tc ase* is expressed in the head appendages (arrows) and in a few cells in the lateral body wall (arrowheads). (B) The latter expression persists into NS11 (small arrowheads). Additional cells express *Tc ase* in the appendages (arrows) and the lateral body wall (large arrowhead). (C) At NS7, *Tc pros* is expressed in the mandibular appendages in addition to the CNS. (D) *Tc sna* expression is visible in the same domains as *Tc pros* in the developing mandibles (arrow) at NS7. In addition, *Tc sna* positive cell clusters appear in the lateral body wall of t2, t3 and the first couple of abdominal segments (arrowheads). (E) At NS9, this expression has extended up to A9 (arrowheads). In addition, *Tc sna* is expressed in clusters in all appendages (arrows). (F) At NS15.1, *Tc ase* expression has decreased in the PNS except for a few cells in the lateral body wall (arrow) and a domain at the tip of the antennae (arrow). (G) At NS10, Tc pros is expressed in clusters of cells in the lateral body wall of t2, t3 and a1-a3. Abbreviations see Suppl. Fig. 1. Scale bar in A, 50 µm in A-E, scale bar in F, 100 µm, scale bar in G, 50 µm.

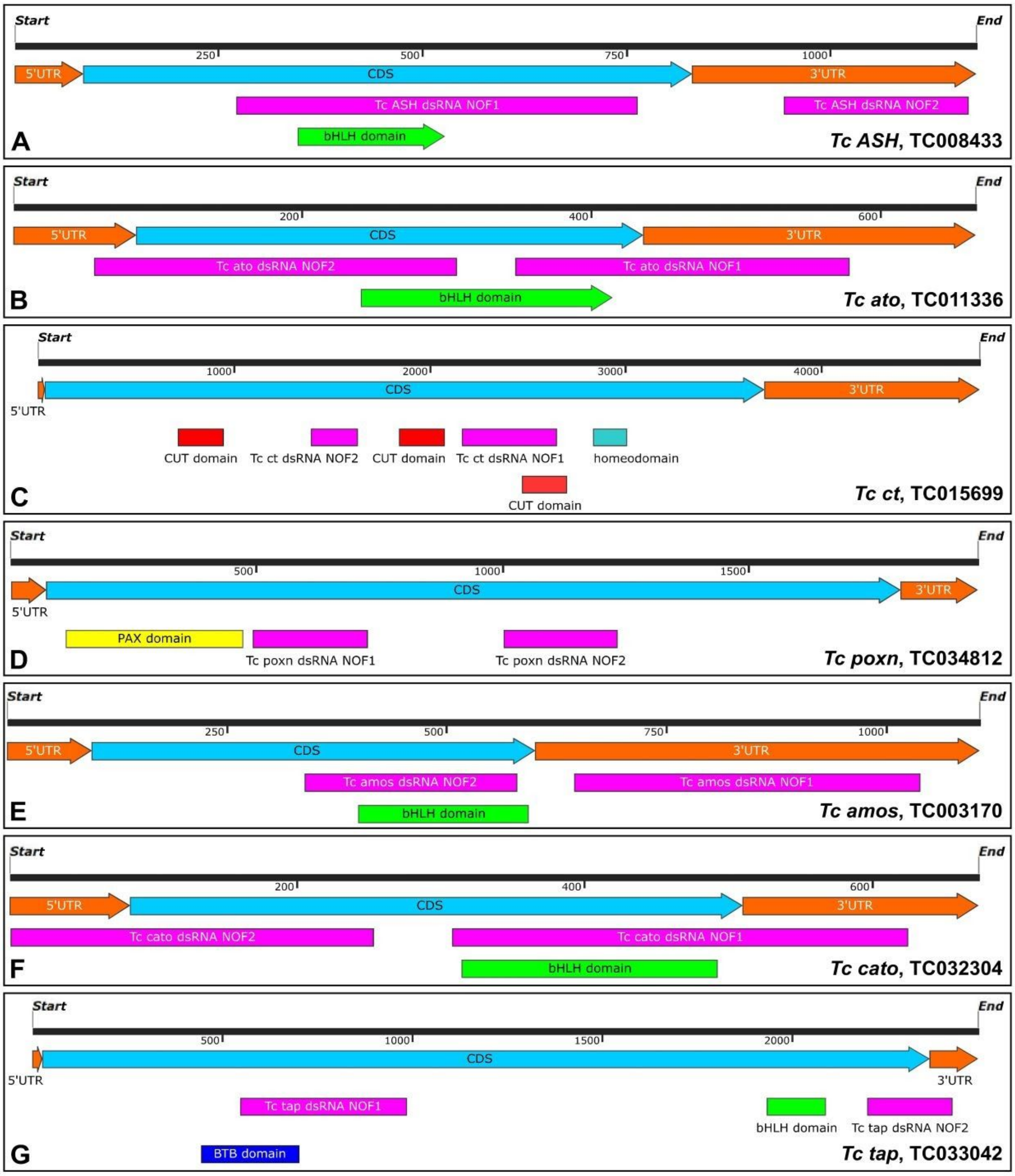


**Supplementary Figure 5. Maps of mRNA architecture of analysed genes**. (A-G) Each panel shows 5’ and 3’UTR (orange) and coding sequences (blue), as well as location of dsRNA NOFs (pink), and the location of the conserved DNA binding domains (in green (bHLH), red (CUT domain), turquoise (homeodomain) and yellow (Pax domain). Gene names and corresponding TC sequence numbers are given in the bottom right corner of each panel. Maps were generated with the SnapGene software (from Insightful Science; available at snapgene.com).

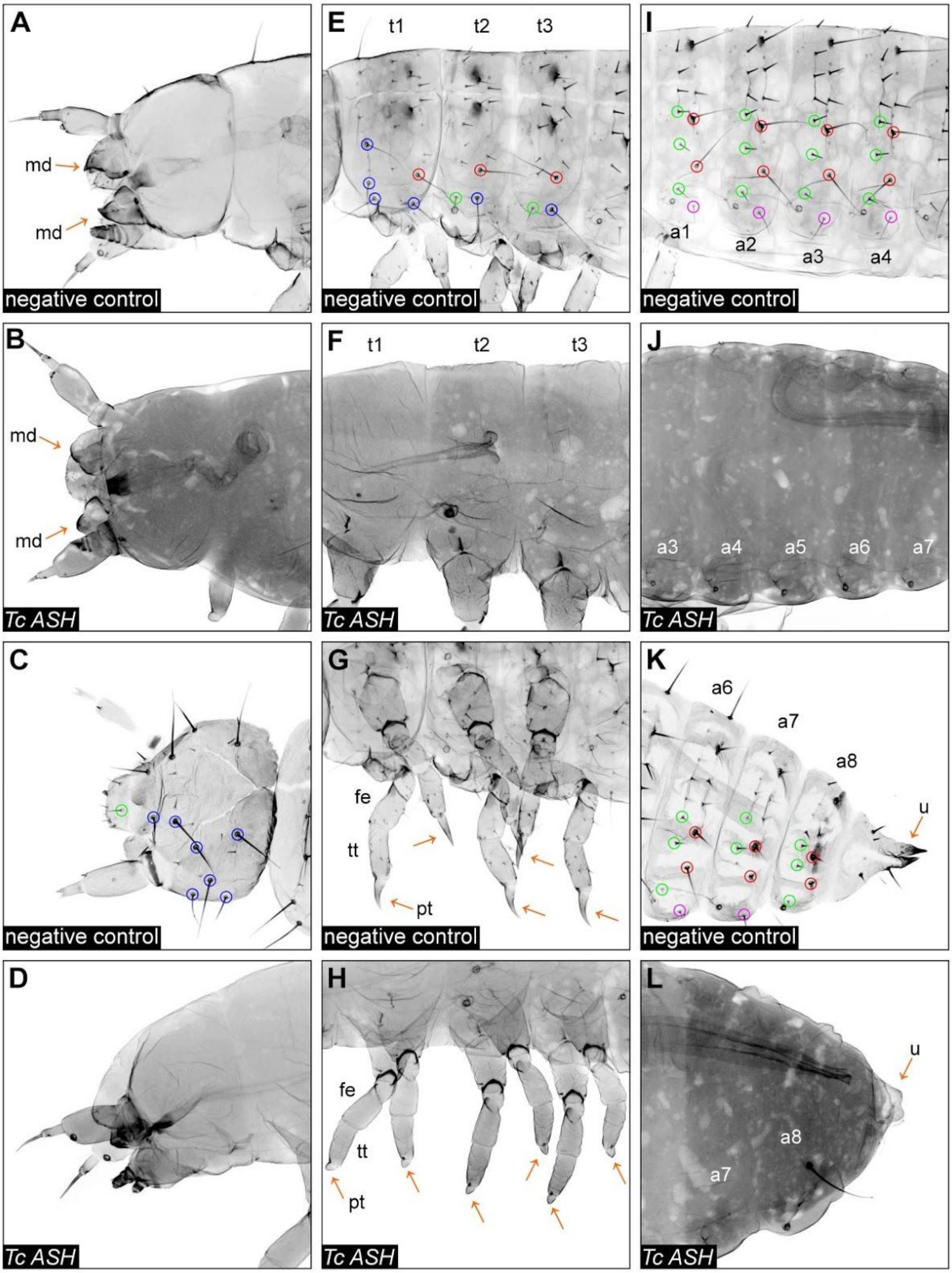


**Supplementary Figure 6 *Tc ASH* RNAi phenotypes.** Confocal images of larval cuticles 1st larval stage). Anterior is to the right. (A, C, E, G, I, K) Negative control cuticles. Please note that the sensilla of the head capsule and thorax are out of focus. (B, D, F, H, J, L) *Tc ASH* RNAi cuticles. (B, F, J, D) The strongest cuticle phenotype resulting from *Tc ASH* RNAi are ‘naked’ larvae, i.e. larvae missing all external sensilla (compare to C, E, I). (B, H, L) *Tc ASH* RNAi also leads to rounded pretarsi (pt) (47.89% (n=263) have at least one pretarsal segment rounded), urogomphi (u) (100% (n=263) have rounded u), and mandibles (md) (83% (n=263) have rounded md) (orange arrows; compare to (A, G, K)). Scale bar, 50 µm.

### Supplementary Tables

**Supplementary Table 1 Double-stranded RNA information.** TC numbers are listed for each gene, as well as the corresponding number from the iBeetle screen (iB number) for each fragment (Schmitt-Engel et al., 2015), including the concentration used for each fragment.

| gene name | ID | TC number | concentration |
|---|---|---|---|
| *Tc ASH* NOF1 | iB_04489 | TC008433 | 1µg/µl |
| *Tc ASH* NOF2 | iB_04489-2 | | 1µg/µl |
| *Tc ato* NOF1 | iB_09565 | TC011336 | 1µg/µl |
| *Tc ato* NOF2 | iB_09565-2 | | 1µg/µl |
| *Tc ct* NOF1 | iB_06353 | TC015699 | 3µg/µl |
| *Tc ct* NOF2 | iB_06353-2 | | 3µg/µl |
| *Tc poxn* NOF1 | iB_10412 | TC034812 | 1µg/µl |
| *Tc poxn* NOF2 | iB_10412-2 | | 1µg/µl |
| *Tc cato* NOF1 | XX-90422-1* | TC032304 | 1µg/µl |
| *Tc cato* NOF2 | XX-90422-2* | | 1µg/µl |
| *Tc tap* NOF1 | iB_02207 | TC033042 | 1µg/µl |
| *Tc tap* NOF2 | iB_02207-2 | | 1µg/µl |
| *Tc amos* NOF1 | iB_06876 | TC003170 | 1µg/µl |
| *Tc amos* NOF2 | iB_06876-2 | | 1µg/µl |

* Note that this gene was not analysed in the iBeetle screen (Schmitt-Engel et al., 2015)

**Supplementary Table 2. Summary of RNAi injection results for both non-overlapping dsRNA fragments.** We subdivided the larvae resulting from the RNAi experiments into the following categories: In the control larvae, (1) ‘wt’ includes all larvae that have 98 to 100% sensilla at the analysed positions. The variations in the number of sensilla in specific positions of control larvae are shown in Suppl. Tab. 3; (2) ‘Phenotype’ includes all larvae showing recurring specific sensilla and/or other morphological phenotypes corresponding to the injected double-stranded RNA. (3) ‘Non-specific’ includes all larvae showing sporadic morphological defects, and (4) ‘empty eggs’ includes all eggs that do not develop cuticles. *Tc ASH* RNA*i*: When comparing our results to the iBeetle screen database, we find similar phenotypes for *Tc ASH* NOF1 (targets CDS and overlaps with bHLH domain), and also for NOF2 (targets 3’UTR). We obtained the same range of phenotypes from the injections of both fragments; however, the phenotypes resulting from injections of NOF2 are less strong compared to NOF1. *Tc ato* RNAi: Information on the phenotype for *Tc ato* RNAi from the iBeetle base state that 80% of the injected pupae or adults died 11 days after injection. No other specific phenotype is described. However, we found a specific sensilla phenotype (missing ant_TSOs) for both *Tc ato* RNAi NOF1 (targeting the CDS containing part of the bHLH domain and the 3’UTR) and NOF2 (targeting the 5’UTR and the CDS containing another part of bHLH domain). For NOF1 the penetrance is higher (100% have left and right ant_TSOs missing), while for NOF2 the RNAi effect is milder (41% have left and right ant_TSO missing, 52% have only either left or right ant_TSO missing, 7% have both ant_TSO intact). *Tc ct* RNAi: We initially injected *Tc ct* dsRNA (NOF1 targeting the CDS region, and NOF2 targeting CDS which contains one of the CUT domains) into female pupae. We observed a high lethality (92% for NOF1 and 84% for NOF2, 11 days after injection), as well as sterility as no eggs could be collected. In the iBeetle base the results for *Tc ct* RNAi correspond to our observations, also reporting high female lethality (50% of injected pupae are dead 11 days after injection). To circumvent the problem of lethality, we performed embryonic RNAi and obtained larval sensilla phenotypes for both NOFs. *Tc poxn* RNAi: In the iBeetle base several unspecific phenotypes showing less than 30% penetrance are recorded and a lethality of injected pupae or adults of 30% 11 days after injection was observed. In our injections of NOF1 (targeting the CDS), the lethality falls into a similar range (30% of pupae or adults died 11 days after injection). In contrast to the iBeetle screen results, we found a specific sensilla duplication phenotype. However, we did not observe a phenotype for NOF2 (targeting a different part of the CDS), despite repeating the injections.

| | **negative control pRNAi** | | | | | **negative control eRNAi** | | | | |
|---|---|---|---|---|---|---|---|---|---|---|
| | wt | phenotype | non-specific | empty eggs | total | wt | phenotype | non-specific | empty eggs | total |
| ∑ | 86 | 0 | 52 | 66 | 204 | 22 | 0 | 4 | 0 | 4 |
| % | 42.16% | 0.00% | 25.49% | 32.35% | | 84.62% | 0.00% | 15.38% | 0.00% | |
| | ***Tc ASH* NOF1** | | | | | ***Tc ASH* NOF2** | | | | |
| | wt | phenotype | non-specific | empty eggs | total | wt | phenotype | non-specific | empty eggs | total |
| ∑ | 0 | 91 | 5 | 45 | 141 | 0 | 172 | 20 | 54 | 246 |
| % | 0.00% | 64.54% | 3.55% | 31.91% | | 0.00% | 69.92% | 8.13% | 21.95% | |
| | ***Tc ato* NOF1** | | | | | ***Tc ato* NOF2** | | | | |
| | wt | phenotype | non-specific | empty eggs | total | wt | phenotype | non-specific | empty eggs | total |
| ∑ | 0 | 34 | 17 | 60 | 111 | 25 | 27 | 6 | 50 | 108 |
| % | 0.00% | 30.63% | 15.32% | 54.05% | | 23.15% | 25.00% | 5.56% | 46.30% | |
| | ***Tc ct* NOF1** | | | | | ***Tc ct* NOF2** | | | | |
| | wt | phenotype | non-specific | empty eggs | total | wt | phenotype | non-specific | empty eggs | total |
| ∑ | 1 | 13 | 1 | - | 15 | 0 | 4 | 7 | - | 11 |
| % | 6.67% | 86.67% | 6.67% | - | | 0.00% | 36.36% | 63.64% | - | |
| | ***Tc poxn* NOF1** | | | | | ***Tc poxn* NOF2** | | | | |
| | wt | phenotype | non-specific | empty eggs | total | wt | phenotype | non-specific | empty eggs | total |
| ∑ | 4 | 57 | 14 | 36 | 111 | 63 | 0 | 22 | 42 | |
| % | 3.60% | 51.35% | 12.61% | 32.43% | | 49.61% | 0.00% | 17.32% | 33.07% | |

**Supplementary Table 3 The negative control shows variations in the number of sensilla at specific positions.** Overall 98 to 100% of the sensilla analysed are present in control larvae. The observed variation is due to the absence of sensilla at specific positions along the anterior-posterior axis. In the head and thorax, 99.1% of the analysed sensilla are present at all positions (2,484/2,507), while in the abdominal segments overall 2.87% (199/6912) of sensilla are missing. The abdominal TSOs (4.32%) show the highest variability followed by the CSGs (2.60%).

| | head | | | | |
|---|---|---|---|---|---|
| | n sensilla for 108 larvae | counted | | % | |
| | | wt | wt* | wt | wt* |
| BSMs | 108 | 108 | 0 | 100.00% | 0.00% |
| CSMs | 1080 | 1061 | 19 | 98.24% | 1.76% |
| CSGs | - | | | - | - |
| TSOs | 108 | 106 | 2 | 98.15% | 1.85% |
| | **thorax** | | | | |
| | n sensilla for 108 larvae | counted | | % | |
| | | wt | wt* | wt | wt* |
| BSMs | 216 | 215 | 1 | 99.54% | 0.46% |
| CSMs | 648 | 648 | 0 | 100.00% | 0.00% |
| CSGs | 324 | 323 | 1 | 99.69% | 0.31% |
| TSOs | - | - | - | - | - |
| | **abdomen** | | | | |
| | n sensilla for 108 larvae | counted | | % | |
| | | wt | wt* | wt | wt* |
| BSMs | 2592 | 2550 | 42 | 98.38% | 1.62% |
| CSMs | - | - | - | - | - |
| CSGs | 1728 | 1683 | 45 | 97.40% | 2.60% |
| TSOs | 2592 | 2480 | 112 | 95.68% | 4.32% |

**Supplementary Table 4** ***Tc ASH*** **RNAi quantification of phenotypes.** The table shows the break-down of the phenotype by sensilla category and body section (head, thorax, abdomen) All larvae that showed a phenotype (i.e. missing sensilla) were included in the analysis (The table shows the break-down of the phenotype by sensilla category and body section (head, thorax, abdomen). All larvae that showed a phenotype (i.e. missing sensilla) were included in the analysis (NOF1 (n = 91) plus NOF2 (n = 172)).

| | **head** | | | | |
|---|---|---|---|---|---|
| | **n sensilla for 263 larvae** | **counted** | | **%** | |
| | | **wt** | **phenotype** | **wt** | **phenotype** |
| **BSMs** | 263 | 1 | 262 | 0.38% | 99.62% |
| **CSMs** | 2630 | 253 | 2377 | 9.62% | 90.38% |
| **CSGs** | - | - | - | - | - |
| **TSOs** | 263 | 256 | 7 | 97.34% | 2.73% |
| | **thorax** | | | | |
| | **n sensilla for 263 larvae** | **counted** | | **%** | |
| | | **wt** | **phenotype** | **wt** | **phenotype** |
| **BSMs** | 526 | 103 | 423 | 19.58% | 80.42% |
| **CSMs** | 1578 | 765 | 813 | 48.48% | 51.52% |
| **CSGs** | 789 | 488 | 301 | 61.85% | 38.15% |
| **TSOs** | - | - | - | - | - |
| | **abdomen** | | | | |
| | **n sensilla for 263 larvae** | **counted** | | **%** | |
| | | **wt** | **phenotype** | **wt** | **phenotype** |
| **BSMs** | 6312 | 2570 | 3742 | 40.72% | 59.28% |
| **CSMs** | - | - | - | - | - |
| **CSGs** | 4208 | 2186 | 2022 | 51.95% | 48.05% |
| **TSOs** | 6312 | 1654 | 4658 | 26.20% | 73.80% |

**Supplementary Table 5 *Tc ato* RNAi quantification of phenotypes.** The table shows the break-down of the phenotype by sensilla category and body section (head, thorax, abdomen). All larvae that showed a phenotype (i.e. missing sensilla, reduced sensilla length) were included in the analysis (NOF1 (n = 34) and NOF2 (n = 27)).

| | head | | | | |
|---|---|---|---|---|---|
| | n sensilla for 61 larvae | counted | | % | |
| | | wt | phenotype | wt | phenotype |
| BSMs | 61 | 61 | 0 | 100.00% | 0.00% |
| CSMs | 610 | 577 | 33 | 94.59% | 5.41% |
| CSGs | - | | | - | - |
| TSOs | 61 | 2 | 59 | 3.28% | 96.72% |
| | **thorax** | | | | |
| | **n sensilla for 61 larvae** | **counted** | | **%** | |
| | | **wt** | **phenotype** | **wt** | **phenotype** |
| BSMs | 122 | 115 | 7 | 94Z26% | 5.74% |
| CSMs | 366 | 354 | 12 | 96.72% | 3.28% |
| CSGs | 183 | 181 | 2 | 98.91% | 1.09% |
| TSOs | - | - | - | - | - |
| | **abdomen** | | | | |
| | **n sensilla for 61 larvae** | **counted** | | **%** | |
| | | **wt** | **phenotype** | **wt** | **phenotype** |
| BSMs | 1464 | 1380 | 84 | 94.26% | 5.74% |
| CSMs | - | - | - | - | - |
| CSGs | 976 | 951 | 25 | 97.44% | 2.56% |
| TSOs | 1464 | 1367 | 97 | 93.37% | 6.63% |

**Supplementary Table 6 *Tc ct* RNAi quantification of phenotypes.** The table shows the break-down of the phenotype by sensilla category and body section (head, thorax, abdomen). All larvae that showed a phenotype (i.e. missing sensilla, reduced sensilla length, socket only) were included in the analysis (NOF1 (n = 13) and NOF2 (n = 4)).

| | **head** | | | | |
|---|---|---|---|---|---|
| | **n sensilla for 17 larvae** | **counted** | | **%** | |
| | | **wt** | **phenotype** | **wt** | **phenotype** |
| **BSMs** | 17 | 11 | 6 | 64.71% | 35.29% |
| **CSMs** | 170 | 71 | 99 | 41.76% | 58.24% |
| **CSGs** | - | - | - | - | - |
| **TSOs** | 17 | 1 | 16 | 5,.8% | 94.12% |
| | **thorax** | | | | |
| | **n sensilla for 17 larvae** | **counted** | | **%** | |
| | | **wt** | **phenotype** | **wt** | **phenotype** |
| **BSMs** | 34 | 22 | 12 | 64.71% | 35.29% |
| **CSMs** | 102 | 50 | 52 | 49.02% | 50.98% |
| **CSGs** | 51 | 24 | 27 | 47.06% | 52.94% |
| **TSOs** | - | - | - | - | - |
| | **abdomen** | | | | |
| | **n sensilla for 17 larvae** | **counted** | | **%** | |
| | | **wt** | **phenotype** | **wt** | **phenotype** |
| **BSMs** | 408 | 231 | 176 | 56.62% | 43.14% |
| **CSMs** | - | - | - | - | - |
| **CSGs** | 272 | 129 | 143 | 47.43% | 52.57% |
| **TSOs** | 408 | 46 | 364 | 11.27% | 89.22% |

**Supplementary Table 7 *Tc poxn* RNAi quantification of phenotypes.** The table shows the break-down of the phenotype by sensilla category and body section (head, thorax, abdomen). All larvae that showed a phenotype (i.e. missing sensilla, duplicated sensilla) were included in the analysis (NOF1 (n = 57), NOF2 cuticles showed no phenotype).

| | **head** | | | | |
|---|---|---|---|---|---|
| | **n sensilla for 57 larvae** | **counted** | | **%** | |
| | | **wt** | **phenotype** | **wt** | **phenotype** |
| **BSMs** | 57 | 57 | 0 | 100.00% | 0.00% |
| **CSMs** | 570 | 562 | 8 | 98.60% | 1.40% |
| **CSGs** | - | - | - | - | - |
| **TSOs** | 57 | 47 | 10 | 82.46% | 17.54% |
| | **thorax** | | | | |
| | **n sensilla for 57 larvae** | **counted** | | **%** | |
| | | **wt** | **phenotype** | **wt** | **phenotype** |
| **BSMs** | 114 | 107 | 7 | 93.86% | 6.14% |
| **CSMs** | 342 | 249 | 93 | 72.81% | 27.19% |
| **CSGs** | 171 | 171 | 0 | 100.00% | 0.00% |
| **TSOs** | - | - | - | - | - |
| | **abdomen** | | | | |
| | **n sensilla for 57 larvae** | **counted** | | **%** | |
| | | **wt** | **phenotype** | **wt** | **phenotype** |
| **BSMs** | 1368 | 1027 | 341 | 75.07% | 24.93% |
| **CSMs** | - | - | - | - | - |
| **CSGs** | 912 | 894 | 18 | 98.03% | 1.97% |
| **TSOs** | 1368 | 925 | 443 | 67.62% | 32.38% |
